# Herbivores and pathogens can modulate plant population responses to future climate conditions

**DOI:** 10.64898/2026.07.07.736959

**Authors:** Martin Andrzejak, Tiffany M. Knight, Lotte Korell

## Abstract

Climate change is expected to alter plant populations not only through direct environmental shifts but also via changes in biotic interactions, such as with herbivores and pathogens. As plant species are also expected to differ in their responses to both climate and antagonists, plant responses to both factors are expected to be variable and species-specific. To assess whether interactive effects of climate and antagonists on plant population dynamics are common and whether the strength and direction of plant responses vary across species, we conducted a multi-year field experiment that manipulated realistic climate change and experimentally reduced insect herbivores and fungal pathogens. We measured responses of plant vital rates, such as survivorship, growth, and reproduction across six grassland species. Using Integral Projection Models (IPMs) and Life Table Response Experiments (LTREs), we quantified changes in population growth rate across experimental treatments and the contribution of each vital rate to that observed change. Two of the study species declined so drastically over the course of the experiment that demographic quantification of population growth rates was not possible. From the remaining species, *Bromus erectus* and *Plantago lanceolata* show significant interactive responses of climate and antagonist reduction on population growth rates. In contrast, *Dianthus carthusianorum* and *Tragopogon orientalis* showed limited responses to experimental treatments. Notably, our results indicate that in some species biotic interactions may amplify the effects of climate change: the presence of plant antagonists exacerbates the negative effects of the future climate treatment on plant population dynamics. Our findings highlight the complexity in predicting plant population responses to climate change and provide insights for grassland management under future environmental conditions.

## Introduction

Global change is acting on plant populations in a multivariate manner, e.g. directly via changing the abiotic resources for plants but also indirectly via changes in biotic interactions (e.g., herbivory, pathogen infection). Demographic studies provide insights about how underlying changes in survival, growth and reproduction will combine to influence plant population growth rates (Caswell, 2001). Due to the labor required to manipulate direct and indirect effects of climate change and measure demographic vital rates across the life cycle of a plant species, it is common for demographic studies to manipulate a single factor, to concentrate on one vital rate (e.g., growth), or to measure responses for a single species (Ehrlén et al., 2016; García and Ehrlén, 2002; Knight, 2004; Maron and Crone, 2006).

However, we expect that multiple factors will interactively influence population growth (Morris et al., 2020). For example, abiotic climate factors such as temperature and precipitation are expected to directly influence survival and growth of plants (Midolo and Wellstein, 2020), but these abiotic factors might also indirectly influence plant vital rates by altering the strength of biotic interactions (e.g. interactions with pathogens and insect herbivores) (Surówka et al., 2020). Further, direct and indirect effects of climate factors are expected to vary across species with different traits and life history strategies (Boyko et al., 2023; Henn et al., 2018), and understanding these species-specific responses is necessary to scale predictions from populations to communities. Additionally, plant responses to environmental change are often constrained by trade-offs in resource allocation that can only be quantified if multiple vital rates are measured. In particular, plants must balance investment in growth with investment in defense against natural enemies (Herms and Mattson, 1992). Under changing climatic conditions, shifts in resource availability or physiological stress may therefore alter this balance, potentially constraining plant defense responses to pathogens and herbivores (He et al., 2022).

Climate change has strong direct effects on the population dynamics of plant species and shapes the composition of grasslands (Ehrlén, 2019; Gibson and Newman, 2019; Girdler et al., 2025; Hobbs et al., 2007; Ickin et al., 2025) through multidimensional changes in surface temperature and precipitation patterns (Calvin et al., 2023). In central Europe, climate models project warmer temperatures and a decrease in summer precipitation, while spring and fall precipitation is projected to increase (Rockel et al., 2008; Steensen et al., 2026; Wagner et al., 2013). Especially the drier summers can be harmful to plant populations as drought events can cause stress to plants (Hemati et al., 2022) and the plant stress hypothesis states that stressed plants (e.g., exposed to drought) are less able to invest in defenses against plant antagonists (Hamann et al., 2021).

It is well known that plant antagonists (like insects and pathogens) can reduce seed set (Goss and Bergelson, 2007; Kempel et al., 2025; Strauss, 1997), negatively influence survival rate of plant individuals (Jarosz and Davelos, 1995) and decrease plant growth (Van Dijk et al., 2021). These decreases in plant fitness at different phases of the life cycle can result in decreases in overall plant population growth (Tylianakis et al., 2008; Wise, 2023; Ehrlén et al., 2016; García and Ehrlén, 2002; Knight, 2004; Maron and Crone, 2006). As expected, most studies that examined the effects of herbivory found at least a small negative effect on plant population growth (e.g. García and Ehrlén, 2002; Knight, 2004; Maron and Crone, 2006). Pathogens can also negatively affect plant population growth rates (Koupilová et al., 2023; Roy et al., 2011), often through reductions in plant reproduction. However, comparatively few demographic studies have examined the effects of pathogens on plant population dynamics. The meta-analysis of Ehrlén et al. (2016) found only two studies related to pathogens and plant population dynamics (Davelos and Jarosz, 2004; Roy et al., 2011).

To date, the interactive effects of pathogens and insect herbivores on plant populations are still poorly understood (Burdon, 1987; Jarosz and Davelos, 1995; Mordecai, 2011; van Dijk et al., 2021). A meta-analysis from Hauser et al. (2013) found that there was no interactive effect of the two groups of antagonists on plant performance. However, another study has shown a synergistic negative impact of the combined effect of insects and pathogens on plant fitness (e.g., due to insect herbivores preferentially feeding on the remaining healthy plants, Rostás et al., 2003). Further, some field studies showed that insect herbivory may be followed by an infection with pathogens leading to exacerbated negative effects of the herbivory on the fitness of the plant species (Dodd, 1942; Hatcher and Paul, 2001).

Climate change potentially alters plant-herbivore-pathogen interactions. A review by Eastburn and colleagues (2011) showed that the plant-pathogen interactions are changing under future climate conditions, by changing the incidence or severity of plant diseases (see also Burdon et al., 2006; Chakraborty, 2005). Other studies focused on the effect of climate change on the plant-herbivore interactions, revealing that those will be altered in the future, harming plant populations but also highlighting that this effect is species-dependent (DeLucia et al., 2012; Hamann et al., 2021). In a literature review, Trębicki et al. (2017) discussed the possibility that climate change may increase the interactive effects of the antagonists on plant populations. One point made by this study was that it might be difficult to disentangle direct effects of climate change on the antagonist’s interactions (e.g. facilitate or reduce insect outbreaks, which in return can be vectors for pathogens) from indirect effects (e.g., via changed biochemistry of plants due to increased temperature). To our knowledge, there are no studies that investigate the separate and combined effects of herbivores and pathogens on populations of multiple plant species and how they are altered by climate change in a field experiment. Most studies quantifying the effects of herbivores and pathogens are conducted without considering the influence of climate change (Hatcher and Paul, 2001), focus on a single species (Heimes et al., 2015) or focus on a single vital rate (Van Dijk et al., 2020).

Going beyond single vital rates, Integral Projection Models (IPMs) are ideal for assessing potential interactive effects of multiple factors on vital rates across the life cycle and on plant population dynamics (Childs et al., 2003; Jacquemyn et al., 2012; Yule et al., 2013).

Sensitivity analyses of IPMs can pinpoint the vital rates that, if changed, have the largest influence on population growth rates and Life Table Response experiments (Caswell, 1989) can decompose which vital rates are most responsible for observed changes in population growth rates across experimental treatments.

Understanding how plant populations respond to climate change (Hobbs et al., 2009; Maalouf et al., 2012), and whether these responses are due to direct effects of climate change or indirect effects through shifts in plant-antagonist interactions (i.e. herbivores and pathogens), is key to restore and maintain biodiversity in future ecosystems. Our study is located in a large experimental platform with a realistic climate change treatment based on regional climate scenarios (Korell et al. 2020, Schädler et al. 2019), the Global Change Experimental Facility (GCEF). Within this large experimental platform, we established an insect and pathogen removal experiment within species rich grasslands of low land use intensity (mowing). The study was targeted at six different plant species, which we expected to differ in the response of their population dynamics to our treatments. In general, we expected negative effects of future climate as well as negative effects of the different antagonists on population growth rate. We further expect interactions between the two antagonist groups, insects and pathogens, as well as between antagonists and experimental climate change. Specifically, we expect synergistic (negative) effects of future climate and insect and pathogen presence on plant population growth rate (e.g., related to altered plant defenses).

## Methods

### Study site

Our experiment was set up within the experimental platform of the Global Change Experimental Facility (GCEF), which was established in 2013 at the field research station in Bad Lauchstädt (51°22060 N, 117 11°50060 E, 118 m a.s.l.). The GCEF is a two-factorial, large experiment comprising two climate treatments (ambient and future climatic conditions), crossed with five land-use treatments (for more details see (Schädler et al., 2019). To contrast ambient and future climate, 10 plots each of 80 m x 24 m area were randomly assigned to one out of the two climate treatments. The future climate treatment is based on the mean of 12 regional climate models (Schädler et al., 2019) and includes manipulation of precipitation and temperature using mobile roof systems. In spring (March - May) and fall (September - November), precipitation is increased by about 10% using sprinkler systems, while in summer (June - August) precipitation is reduced by about 20% by closing the mobile roofs. From spring to fall, roofs are also closed during nighttime to achieve passive warming, leading to a mean daily temperature increase by 0.55 °C at a height of 5 cm and by 0.24 °C at a height of 70 cm (see Schädler et al., 2019). Plots assigned to ambient climate are equipped with similar constructions but without mobile roofs. Each of these 10 plots is subdivided into five smaller plots of 16 m x 24 m randomly assigned to five land-use types, including various farming and grassland systems. For our study, we used diverse, extensively used (i.e, low management intensity) meadows that were sown with mixtures of 56 grassland species covering all functional groups (grasses, legumes, non-legume forbs). The sown species are typical for mesophilous to dry meadows of the region and the seed material of the species stems from one to three regional populations. The extensively managed meadows are mown regularly twice per year (early June and late August/early September). However, in 2018, 2019, 2021 and 2022, the meadows were mown only once in each year due to the extreme drought that occurred in those years and the related slow regrowth of the vegetation.

In 2015, four subplots of 1.5 x 1.5 m were established on each extensively managed meadow plot and randomly assigned to one of the following reduction treatments: i) an insecticide treatment to reduce the impact of insect herbivory (I), ii) a fungicide treatment to reduce fungal pathogens (P), iii) a combination thereof (IP), or iv) only water applied in similar amounts as a control treatment (C). Pesticides were applied five times per year during the growing season (April - September) except for in 2020 when the last application of pesticides was done in November, in 2021 and 2022 with the last application in October. Acantho, later replaced by Ortivia (due to expiring permits), and Score were used as fungicides and applied alternately onto the plots. Calypso, which was later replaced by Karate Zeon (due to expiring permits), were used as insecticide (Table A1). Fungicides and insecticides were individually or in combination diluted in one liter of water (444 mL/m^2^) that was then applied to the plots. Control plots received the same amount of water without any chemicals.

The spatial arrangement of the climate, fungicide and insecticide treatments represented a three-factorial fully crossed split-plot design, with climate manipulated at the main plot level, and fungicide and insecticide at the subplot level. In total, it resulted in the following eight treatment combinations, each with five replicates: (1) ambient - C, (2) ambient - I, (3) ambient - P, (4) ambient - IP, (5) future - C, (6) future - I, (7) future - P, (8) future - IP.

### Study species

In 2018, we chose six grassland species covering different life-histories, functional groups and statures. *Dianthus carthusianorum* L. was selected one year later in 2019. The selected species are a representative subset of another experiment within the GCEF comparing the effect of climate change on population dynamics in meadows and pastures (Andrzejak et al., 2025). During the course of our experiment, two species (*Lotus corniculatus* and *Scabiosa ochroleuca)* went quasi extinct, i.e. they had too few adult and flowering individuals to parametrize an IPMs.

### Demographic data collection

Demographic data collection started in May 2018 and was repeated until 2021, i.e., for three years. Each 1.50 x 1.50 m^2^ antagonist reduction subplot was divided into five 50 x 50 cm^2^ quadrats on which the coordinates of individuals of the selected species were determined to relocate them the year after. Corners of the quadrats were permanently marked with nails and plastic discs. For the demographic models, we require a total sample size per species of at least 50 individuals per treatment combination. Thus, on each subplot we tried to sample at least 10 individuals per species. The size of an individual was either measured as basal area (cm^2^), for which we measured the extent of the individual horizontal and vertical close to the ground, or we counted the number of leaves (see also Baudraz et al., 2025, Table 1). Shortly before each mowing event at the beginning of June and at the end of August, we counted the number of flowers of each flowering individual within the quadrats. To determine the seed set we collected reproductive units such as seed capsules or seed heads, of individuals outside of the quadrat but within the bigger 1.5 x 1.5 m^2^ subplots. After we counted the seeds per seed capsule or head, we returned the seeds to the respective subplots and distributed them randomly. To determine seedling recruitment for each species, we counted the number of seedlings in spring and autumn on each quadrat. Due to the drought years (2018, 2019 and 2020), two of our study species became quasi extinct (Table 1). In our experiment, a species is categorized as quasi extinction if the number of individuals dropped below 25 or fewer than 10 individuals were flowering per treatment combination.

**Table 1.** Description and ecology for each plant species used in the experiment. The information was extracted from: https://wiki.ufz.de/biolflor/index.jsp. The † in the column Quasi extinct signifies a species that had to be dropped from our analysis while * stands for a species that could be used.

| Species | Growth form | Plant family | Life history | Census | Unit of measure | Quasi extinct |
| --- | --- | --- | --- | --- | --- | --- |
| *Bromus erectus* Huds. | Grass | Poaceae | p<sup>1</sup> | 2018 – 2021 | Basal | * |
| *Dianthus carthusianorum* L. | Grass | Caryophyllaceae | p | 2019 – 2021 | Basal | * |
| *Lotus corniculatus* L. | Legume | Fabaceae | p | 2018 – 2020 | Basal | † |
| *Plantago lanceolata* L. | Forb | Plantaginaceae | p | 2018 – 2021 | Number | * |
| *Scabiosa ochroleuca* L. | Forb | Dipsacoideae | p | 2018 – 2021 | Basal | † |
| *Tragopogon orientalis* L. | Forb | Asteraceae | h<sup>2</sup> | 2018 – 2021 | Number | * |

<sup>1</sup>perennial and iteroparous, <sup>2</sup>pluriennial-hapaxanthic

### Life cycle stages and vital rates

We modeled a year-to-year life cycle (April to April) for the species with a discrete stage class for seedlings and a continuous stage class for adult plants (Figure 1). We did not include a seed bank in our life cycle, since the only species of our studied species that builds a long-lasting seed bank is *Plantago lanceolata* (Chen, 2018). Likewise other demographic studies of this species typically do not incorporate the seed bank stage (e.g. www.plantpopnet.com) because most of the seeds germinate immediately. This information was used to calculate the integral projection model (IPM) for all our study species (see following section for more information).

**Figure 1.**
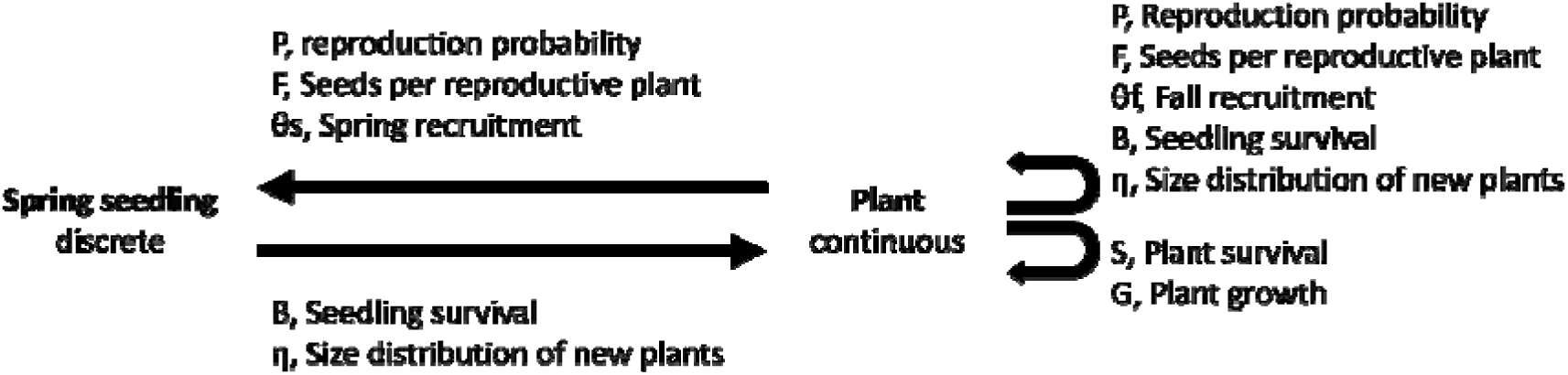
Life cycle graph for each species in our study. Shown is each vital rate that went into the integral projection model (IPM) and the abbreviation used.

As size was a good predictor for the vital rates, we modeled the continuous stages of the IPM against the natural logarithm of the size. Therefore, plant survival (S) is a function of the log transformed size (z) of an individual (i) at t_0_ and if it was still alive at t_1_ (Eq. 2). This function was modeled as a Bernoulli process with probability of survival S_t1_ (Eq. 1). In this model the intercept of the fitted curve is defined as α and the slope is defined as β.

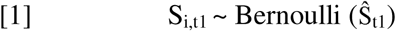

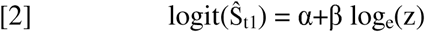

As for plant growth (G), we defined G_i, t1_ as the normally distributed change in z and log transformed size at t1 (z’) of each surviving plant individual (i) (Eq. 3). Growth is modeled as a linear function of z’ for surviving individuals depending on z. Therefore, the intercept is defined as α^G^, the slope as β^G^ (Eq. 4) and the standard deviation as

[3]

[4]

The reproduction probability (P_i, t0_) was modeled as Bernoulli process (Eq. 5). In this kernel we tested how size of an individual (i) at time t_0_ influences the probability of reproduction. In this model the intercept is defined as and the slope as β^P^ using a logit link function (Eq. 6).

[5]

[6]

We calculated the number of seeds (F, Eq. 7) for each reproductive plant. Further we defined the number of seeds produced by an individual (i) at t0 as a function of z, which we modeled as a Poisson distribution. For this vital rate the intercept was defined as and the slope was defined as β^F^(Eq. 8).

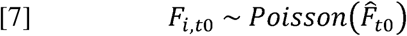

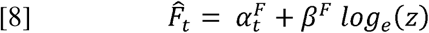

We separated the seedlings that emerged from the total number of seeds (recruitment) into fall (θ*_f, j, t0_*) and spring recruitment (θ_s,_ *_j, t0_*). For that, we calculated the sum of seeds per subplot (S_j,t_, Eq. 9), with S_i,t1_ being defined as the total number of seeds produced by an individual (n) in a subplot at time t_1_.

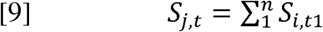

After we calculated this, we divided the result by the number of seedlings in fall t0 (*Rf_j, to_*, Eq. 10) and spring t1 (Rs_j,t1_, Eq. 11), respectively.

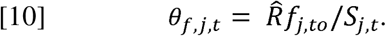

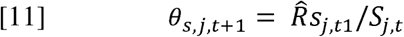

The proportion of seedlings that survived and grew into plants (from t0 to t1) is defined as establishment (B_i_) We calculated this value for each subplot (j). To do so we calculated the total amount of seedlings in spring and fall at time t0 (Rsum_j, t0_) of a subplot. This number was then divided by the number of new individuals at t1 (Ni_j, t1_) in the same subplot (Eq. 12).

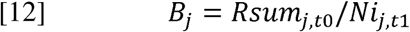

In the last step, we calculated the log size distribution of the new individuals η, which is defined as the normally distributed size of individuals able to enter the continuous plant stage in year t1 (Eq. 13). For that we calculated the mean (log_e_(η_t1_)) and standard deviation (σ_η_) of the size.

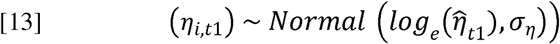

pooled the data across years for each species. This leads to a higher sample size and therefore a higher power to address our questions. The results for the yearly population growth rates can be found in the appendix (Figure A1 – Figure A4).

### Effects of treatments on vital rates

We tested how survival, growth, flowering probability, and number of seeds were influenced by climate, the different antagonist reduction treatments (insect, pathogen, both) and the interaction of climate and the antagonist reduction treatments for each species by using species-specific mixed-effect models and maximum likelihood. To account for the underlying split-plot design of our experiment, we included a random term ‘plot’ nested in climate into the models. Additionally, we ran tests on the discrete vital rates: recruitment, establishment and size distribution of new plants.

### Integral projection model (IPM)

We build an IPM separately for each study species, in each climate treatment and antagonist reduction treatment, calculating a separate population growth rate (λ) for each species and each treatment combination (Eq. 14). We calculated every possible transition from z to z’ in the IPM. We defined the change in number of plants from one year to the other as:

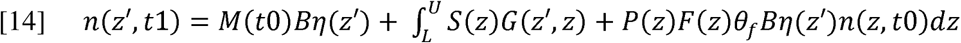

Here the vector n(z’, t1) is defined as the number of plants at size z’ at time t1, with the first term of the function representing the recruitment of spring seedlings (seedlings found in spring entering the adult stage class). This is generally based on the number of spring seedlings at t0, M(t0), the seedling establishment, B, and the size distribution of new plants η(z’). The surface (or the second kernel) describes growth and the survival of plants from t0, n(z, t0) to t1, n(z’, t1), which is described as an integral with the upper limit U (biggest size observed) and a lower limit L (smallest size observed) for each species. We evaluated the integrals across 200 equally sized bins using the midpoint rule, leading to a 200 x 200 matrix. Within our IPM the recruitment of spring seedlings from one year to the next is defined by:

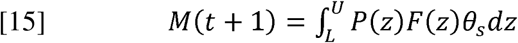

We did not distinguish between new plants from spring seedlings or fall seedlings and we used the same establishment estimate for seedlings (B) twice. By calculating the mean across main effects in the different vital rates (climate and reduction treatments) and afterwards parametrizing a new IPM for each species per main effect. We tested if we could find any main effect of treatments on λ.

### Effects of treatments on population growth rate

We bootstrapped our data a thousand times, calculating lambda for each bootstrap. By doing so we were able to quantify the uncertainty in the estimates of λ. By using a permutation (randomization) test (N = 1000) we were able to establish significance levels between treatments.

### Sensitivity of λ to each parameter

By extracting the parameters for each species and treatment combination from the full statistical model we were able to calculate the sensitivity of λ to these parameters. For that we averaged the parameter for each treatment of interest (for example, averaging the parameters across the ambient: ambient control, ambient insect, ambient pathogen and ambient insect pathogen). After calculating the mean, we permuted each parameter one by one by 0.001 and measured the resulting change in λ by comparing it to the original λ.

### Life table response experiment (LTRE)

By conducting a Life Table Response experiment (LTRE), we calculated the influences of each vital rate on the differences in λ for each pairwise treatment combination. That difference in λ is:

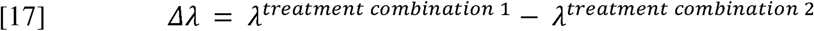

Therefore, the contribution of each vital rate to is:

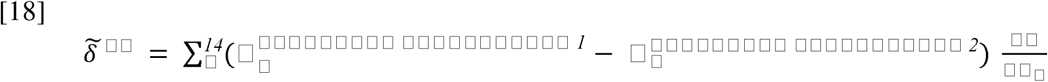

In this function □ is defined as one of the fourteen parameters included in the IPMs while also including the sensitivity of λ to each vital rate, which is defined as 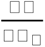 This ensures that vital rates that differ strongly in their magnitude between treatments and vital rates that are much more sensitive to the change of a parameter have a stronger contribution to the LTRE. We combined the LTRE and sensitivity results into five different demographic processes (survival, growth, reproduction, recruitment and establishment, Table 2) to increase understandability.

**Table 2.**
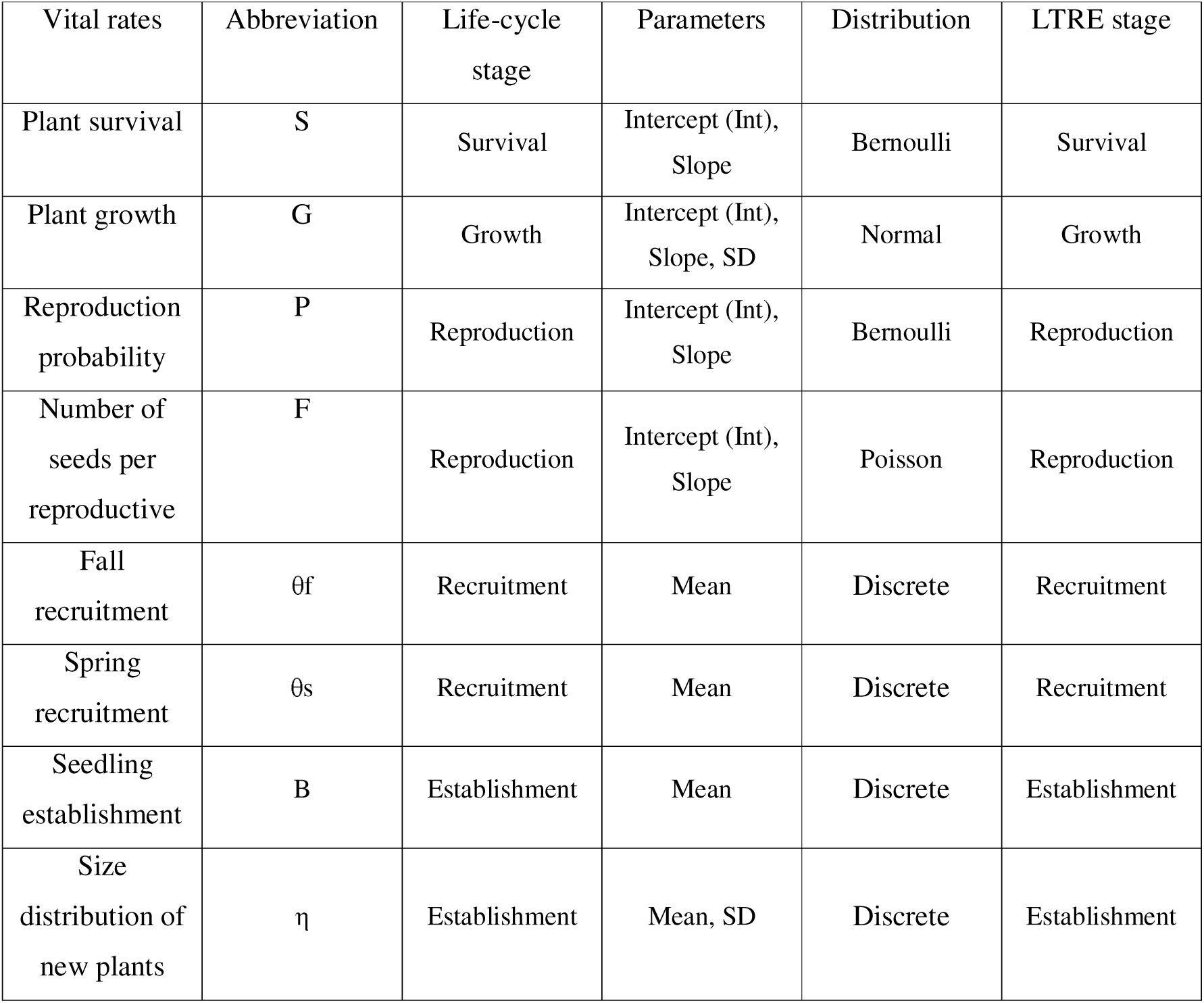
Vital rates and parameters used in the integral projection model (IPM). In the column Distribution discrete indicates vital rates that are not including the continuous stage of size.

| Vital rates | Abbreviation | Life-cycle stage | Parameters | Distribution | LTRE stage |
| --- | --- | --- | --- | --- | --- |
| Plant survival | S | Survival | Intercept (Int), Slope | Bernoulli | Survival |
| Plant growth | G | Growth | Intercept (Int), Slope, SD | Normal | Growth |
| Reproduction probability | P | Reproduction | Intercept (Int), Slope | Bernoulli | Reproduction |
| Number of seeds per reproductive | F | Reproduction | Intercept (Int), Slope | Poisson | Reproduction |
| Fall recruitment | $\theta_f$ | Recruitment | Mean | Discrete | Recruitment |
| Spring recruitment | $\theta_s$ | Recruitment | Mean | Discrete | Recruitment |
| Seedling establishment | B | Establishment | Mean | Discrete | Establishment |
| Size distribution of new plants | $\eta$ | Establishment | Mean, SD | Discrete | Establishment |

### Analysis

We used R (R Core Team, 2022) for all analysis in our study. To calculate the mixed-effect models we used lme4 (Bates et al., 2015), while the IPMs were run using the ipmr package (Levin et al., 2021). The visualization of the results was done with the package ggplot 2 (Wickham, 2016). Further packages used in this study are: openxlsx (Schauberger & Walker, 2021), dplyr (Wickham et al. 2022) and the parallel package (R Core Team, 2022).

## Results

### Effects of climate manipulation on population growth rate

Two of the four species, *Bromus erectus* and *Dianthus carthusianorum,* were projected to increase in population size across both climate treatments, i.e. (log (λ) > 0, Figure 2a and Figure 2b). *Plantago lanceolata* had a population growth rate close to zero under ambient climate conditions but is projected to decline (log () < 0) under future climate conditions. *Tragopogon orientalis* is projected to decline in its population size (log () < 0) in both climate treatments (Figure 2d). In *B. erectus* the population growth rate was significantly higher under ambient compared to future climate conditions (p = 0.001, Figure 2a), while every other species was not significantly influenced by climate.

**Figure 2.**
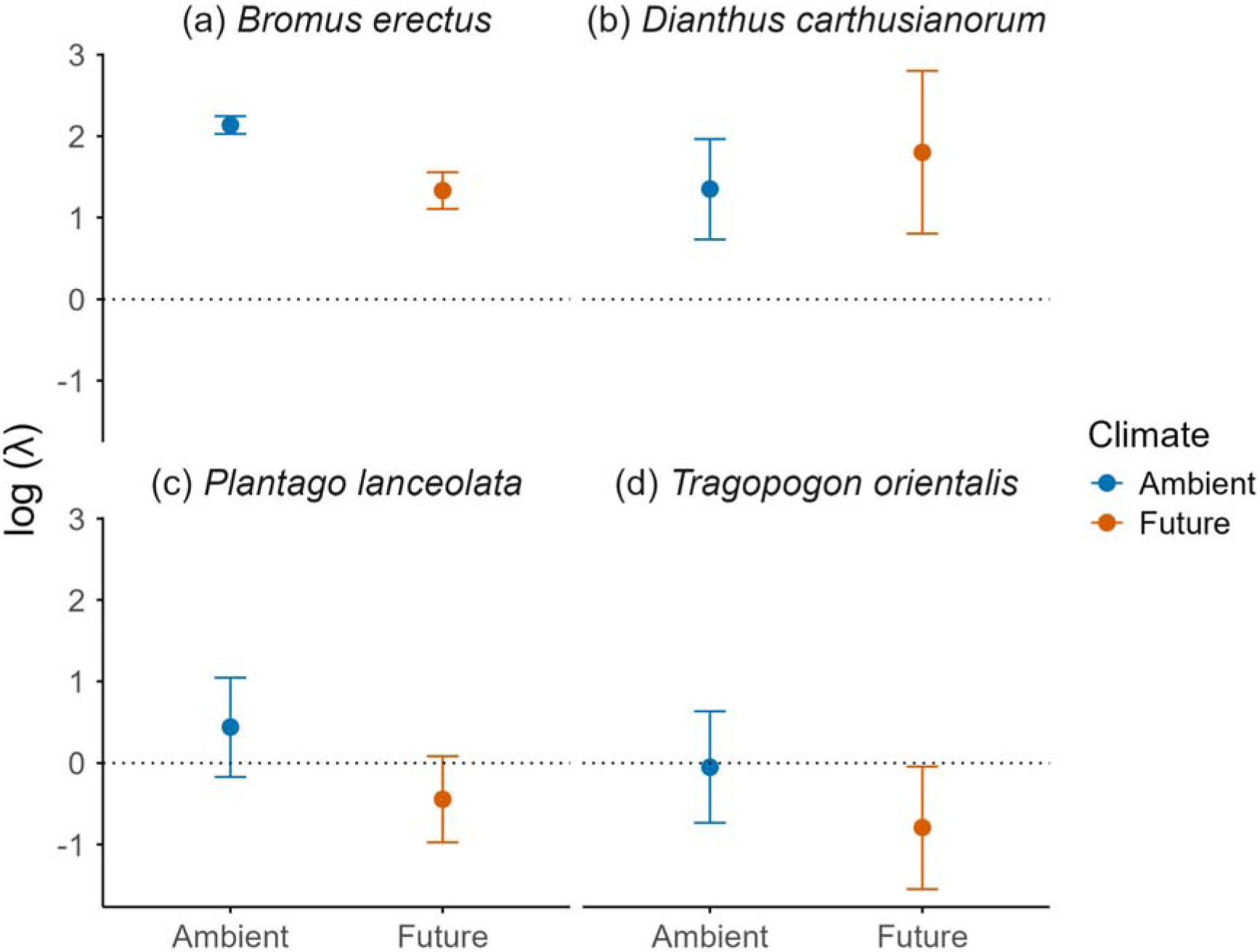
Log population growth rate for (a) *Bromus erectus*, (b) *Dianthus carthusianorum*, (c) *Plantago lanceolata* and (d) *Tragopogon orientalis* for main climate effects (ambient, future). Shown is the mean ± the standard deviation. The horizontal dashed line indicates a neutral population growth rate.

### Effects of antagonist reduction on the population growth rate

*B. erectus* and *D. carthusianorum* both showed a positive population growth rate (log () > 0) across all antagonist treatments (Figure 3a and 3b). Additionally, *B. erectus* had a significantly higher population growth rate in the pathogen reduction treatment compared to the combined exclusion of insect and pathogens (p = 0.046). *P. lanceolata* was projected to decline in the control (C) and pathogen removal (P) (log () < 0), but to increase when both insect herbivores and pathogens are removed (IP log () > 0, Figure 3c). Population growth rate of *P. lanceolata* was higher in I and IP compared to C (p = 0.05 and p = 0.013, respectively, Figure 3c) as well as when I and IP were compared to P (p = 0.034, p = 0.007 respectively, Figure 3c). The population size for *T. orientalis* was projected to decrease in every antagonist reduction treatment and the control (Figure 3d, log (λ) < 0).

**Figure 3.**
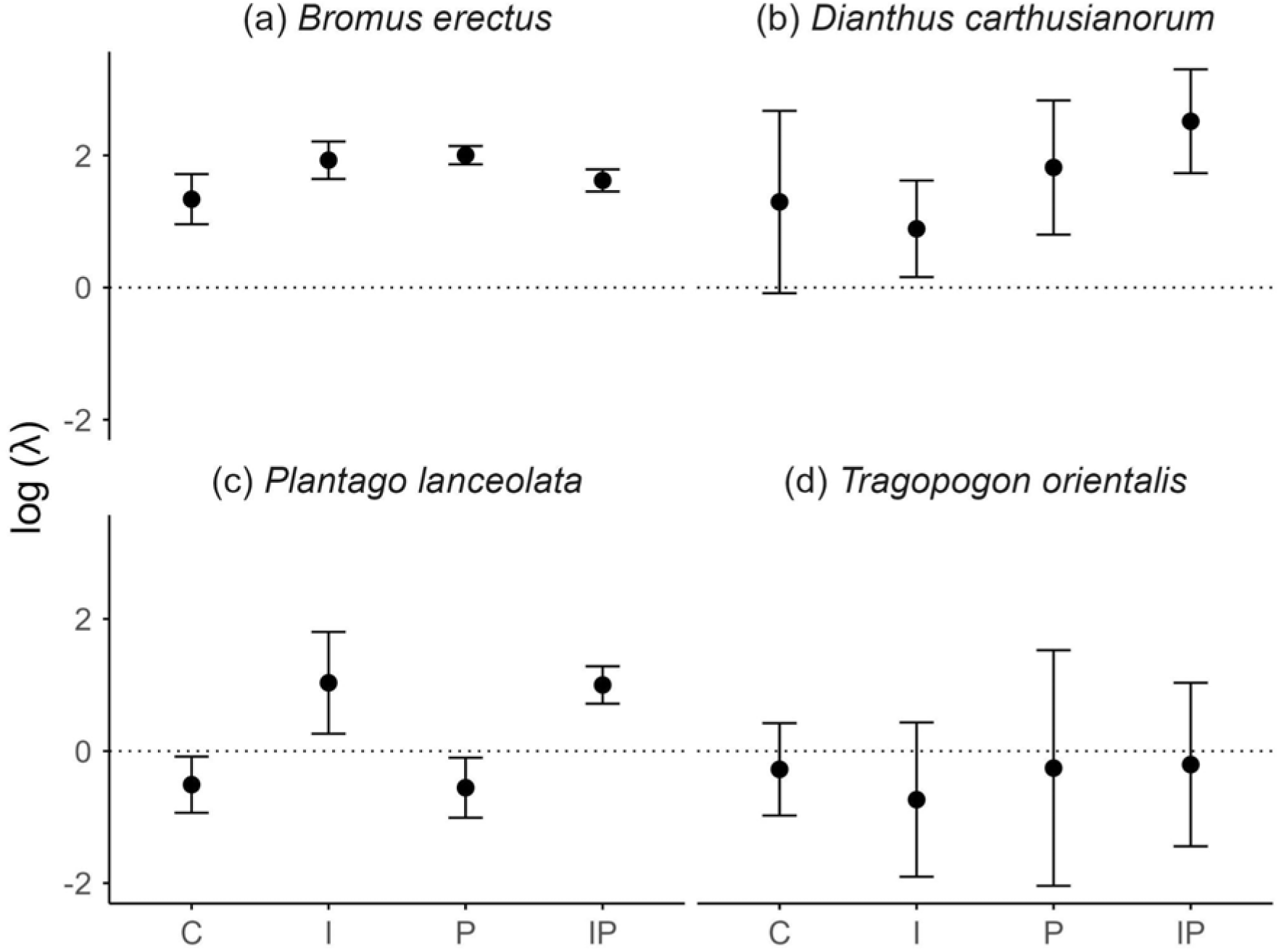
Log of population growth rate (λ) for (a) *Bromus erectus*, (b) *Dianthus carthusianorum*, (c) *Plantago lanceolata* and (d) *Tragopogon orientalis* for the main effect of antagonist reduction (C = control, I = insect, P = pathogen, IP = insect and pathogen). Shown is the mean population growth rate over 1000 bootstrap repetitions and the standard deviation. The horizontal dashed line indicates a neutral population growth rate.

### Interactive effect of climate manipulation and antagonist reduction on the population growth rate

*B. erectus* had a positive population growth rate across all treatment combinations under all experimental conditions, but treatments interactively influenced the magnitude of this projected growth. Under ambient climate conditions, the population growth rate was similar in control and antagonist reduction treatments. In contrast, under future climate conditions, population growth rate was higher when antagonists were removed (Figure 4a). In the future P and I treatments, population growth rate was doubled compared to future C (p = 0.02 and p = 0.039 respectively, Figure 4a). Additionally, population growth of *B. erectus* in the ambient C was significantly higher compared to the future C (p < 0.001, Figure 4a). The same effect could be found between ambient P and future P, where pathogen reduction significantly increased population growth under ambient compared to future conditions (p = 0.011, Figure 4a).

**Figure 4.**
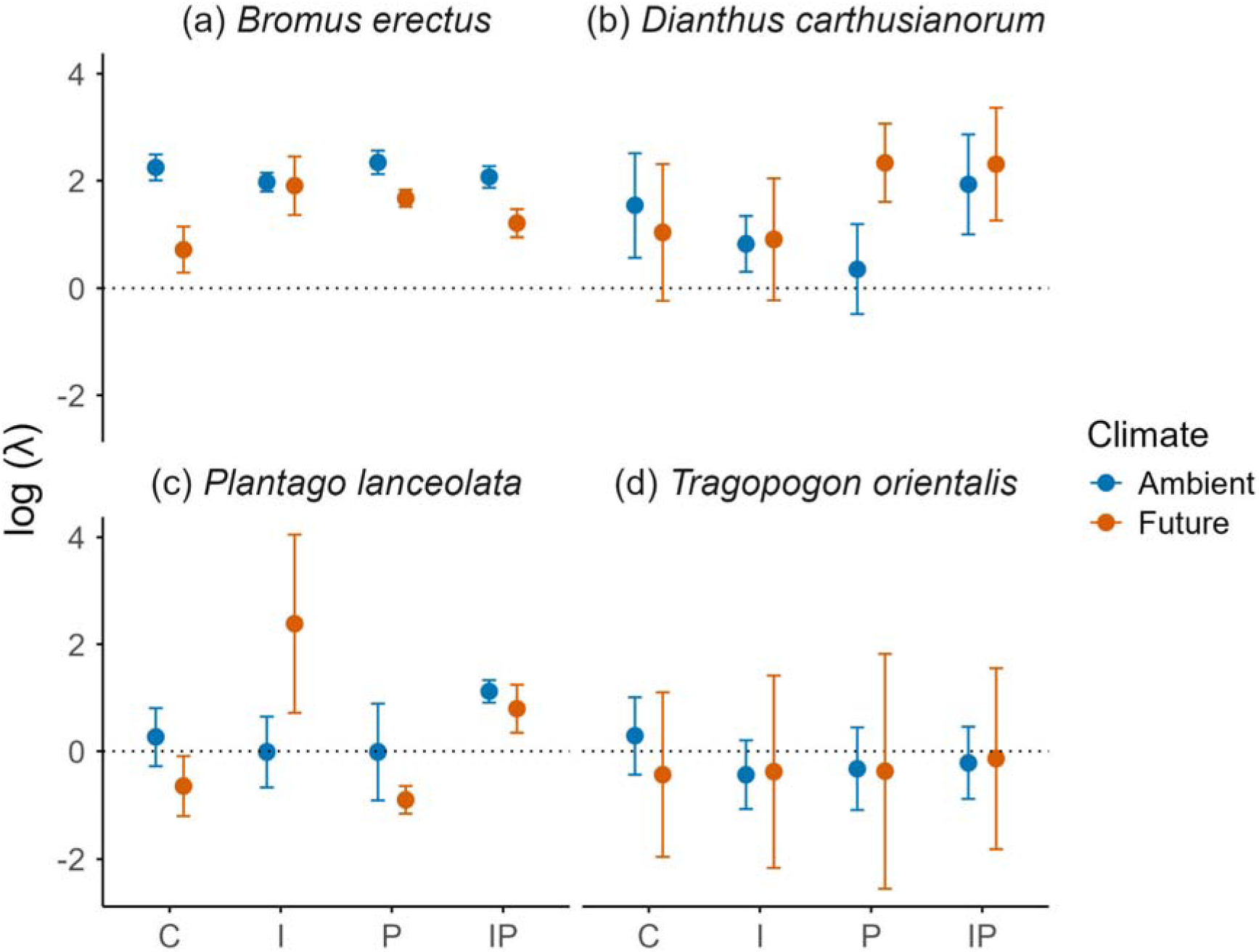
Log population growth rate for (a) *Bromus erectus*, (b) *Dianthus carthusianorum*, (c) *Plantago lanceolata* and (d) *Tragopogon orientalis* for each treatment combination of the two climate treatments and the four reduction treatments (C = control, I = insect, P = pathogen, IP = insect and pathogen). Shown is the mean population growth rate and the standard deviation. The horizontal dashed line indicates a stable population growth rate.

We also found interactive effects of climate manipulation and antagonist reduction for *P. lanceolata*. Under ambient climate conditions, population growth rate of *P. lanceolata* did not differ from zero (i.e. stable population size) in all the reduction treatments, except for the ambient IP treatment where the population growth rate was positive compared to ambient C. Under future climate conditions the effects of antagonist reduction were much more pronounced and either increased (future I and IP log (λ) > 0), or declined (future P (log (λ) < 0), Figure 4c, all p < 0.05) compared to future C.

The population growth rate of *D. carthusianorum* was positive in all treatment combinations with the lowest in the future P treatment (log (λ) > 0). However, population growth rate was higher (marginally significant) in ambient P compared to future P (p = 0.069, Figure 4b).

There was no significant difference in the population growth rate of *T. orientalis* across all treatments (Figure 4d) and the population was decreasing in all treatments except for future C.

### Sensitivity

Across all species and most treatments, the most sensitive vital rates were related to reproduction or recruitment (Figure 5). *T. orientalis* showed high sensitivity towards growth under ambient climate conditions in most treatments, except I under ambient climate conditions. Under future climate conditions, we found a higher sensitivity to changes in survival, as well as vital rates related to individual growth of *T. orientalis* and *P. lanceolata,* across all reduction treatments (Figure 5e – 5h). The sensitivity plots that look only on the main effect climate (Appendix 1, Climate = Figure A5).

**Figure 5.**
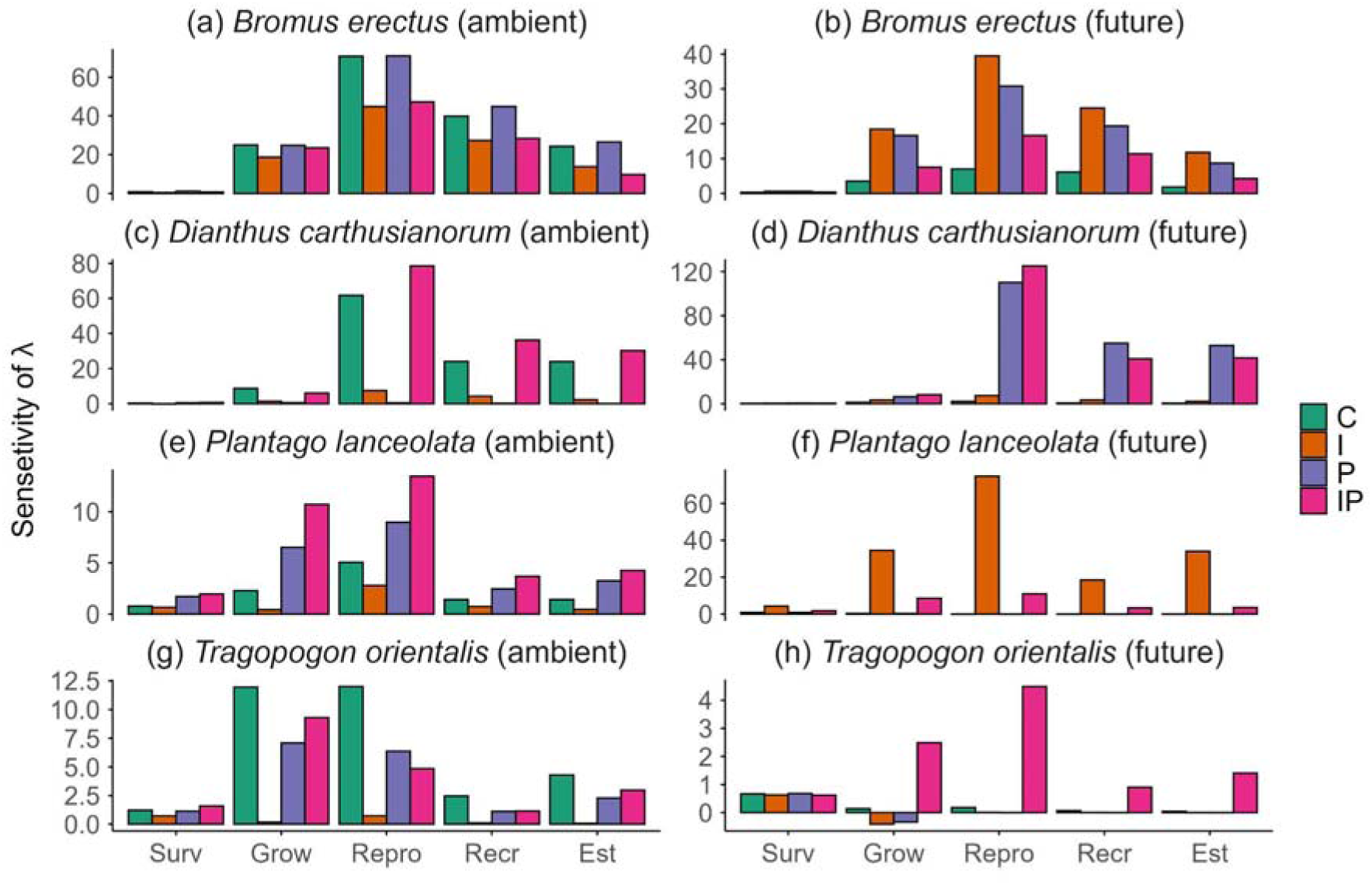
Sensitivity for the pairwise treatment combinations of the reduction under the two different climate treatments. Shown are the different kernels of the Integral projection model. In the figure legend the treatments are abbreviated: C = control, I = insect, P = pathogen, IP = insect and pathogen. Note that, because of visibility, the y-axis of the different plots are not on the same scale. The x-axis abbreviations translate to the following: Est = Establishment, Grow = Growth, Recr = Recruitment, Repro = Reproduction, Surv = Survival. Note that the single plot panels each have a different y-axis scale.

### Life Table Response Experiment

The most dramatic interactive effects of climate and antagonists on log (λ) were seen between the I vs. C treatments in future climate for *B. erectus* and *P. lanceolata* (Figure 4). The LTRE analysis found that changes in reproduction contributed most to the observed change in log (λ) for both species (Figure 6), indicating that under future climate conditions the higher log λ in the I treatment compared to C was due to a higher reproduction. Further for both species the insect removal had beneficial effects on every life stage when compared to the control. The LTRE for the other species and the ambient climate for *B. erectus* and *P. lanceolata* can be found in the supplementary material (Figure A6). The vital rate plots for each life stage and species can also be found in the appendix (Figure A7 - Figure A14)

**Figure 6.**
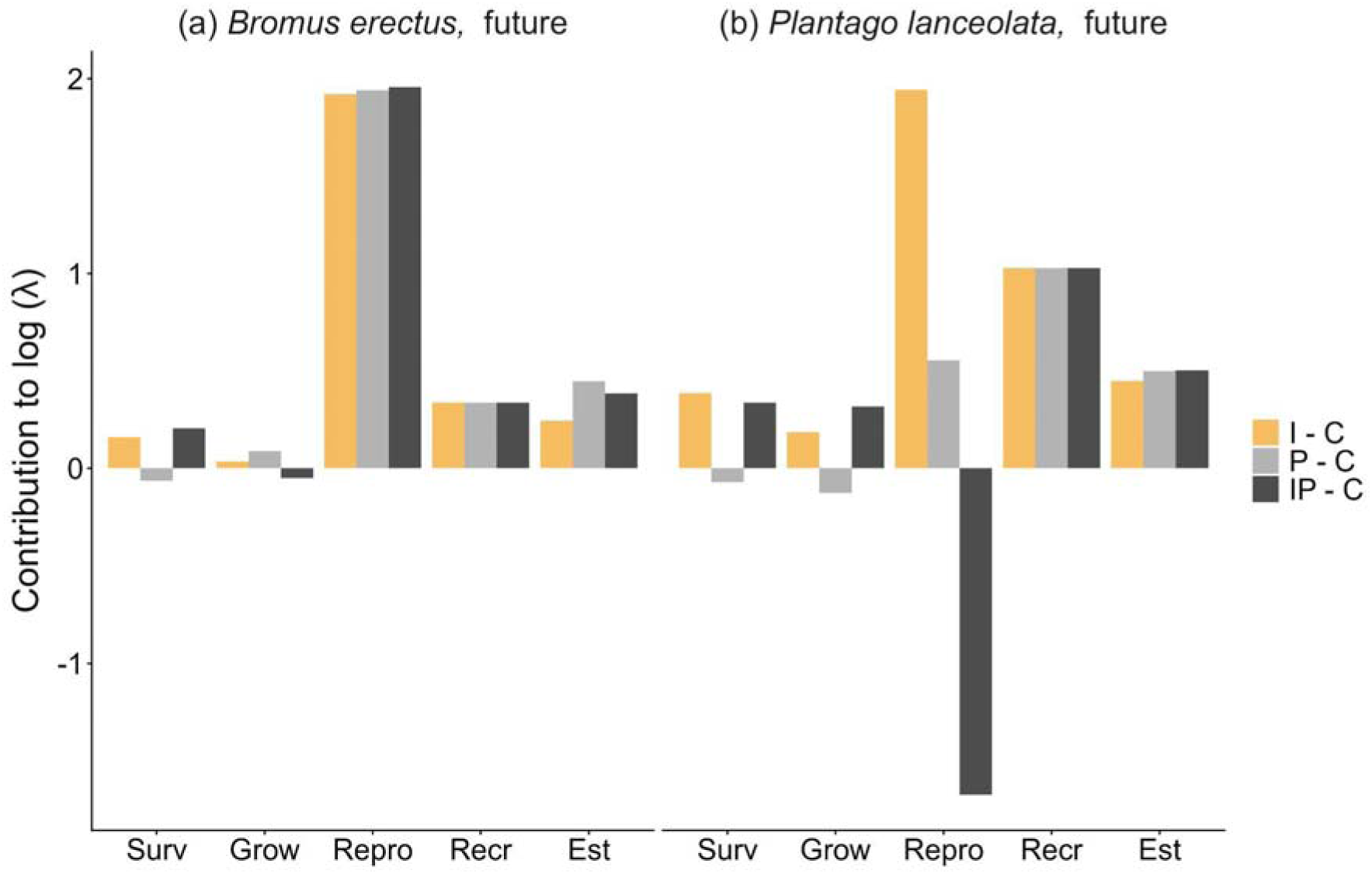
Results of the Life Table Response Experiment (LTRE) for *B. erectus* and *P. lanceolata*. Shown are the results when the reduction treatments are compared to the control under future climate treatments. In the figure legend the treatments are abbreviated: C = control, I = insect, P = pathogen, IP = insect and pathogen. The x axis abbreviations are like this: Est = Establishment, Grow = Growth, Recr = Recruitment, Repro = Reproduction, Surv = Survival. Note that the single plot panels each have a different y-axis scale. The vital rate contributions of the I - C treatment are in orange as they signify the biggest change in log population growth rate.

## Discussion

Across study species we found both interactive and non-interactive effects of climate change and antagonist removal. Two species (*B. erectus, P. lanceolata*) showed stronger responses to antagonist reduction under future climate conditions. This indicates that climate change may indeed alter biotic interactions among plants and antagonists with significant consequences for plant population dynamics.

### Effects of climate and antagonist reduction on population growth rate

*B. erectus,* a grass species that is growing on semi-dry and dry meadows, exhibited a positive population growth rate across all treatment combinations, consistent with previous findings from the same larger experiment, where this species performed well under various management and climate treatments (Andrzejak et al., 2025; Lemmer et al., 2021). Under future but not ambient climate conditions, reduction of insects (I) or pathogens (P) significantly increased population growth rate compared to the control (C). The increased susceptibility of *B. erectus* to insect herbivory and pathogens under future climate conditions supports the plant stress hypothesis (Hamann et al., 2021; Newton et al., 2012; Rhoades, 1983; White, 1969). Given the trade-offs between growth and defense (Herms and Mattson, 1992), resource allocation constraints may further compromise plant responses under future climate conditions (He et al., 2022). Plants usually have to invest resources into defense mechanisms against pathogens and insects, and if those antagonists are reduced (e.g. due to a reduction treatments), plants are able to invest more into growth, and reproduction and still have a high survival rate (Coley et al., 1985; Herms and Mattson, 1992; Simms and Rausher, 1987), thus increasing overall population growth rate. Generally *B. erectus* is predicted to increase in population sizes in the future, threatening the calcareous grassland diversity of central Europe (Poniatowski and Fartmann, 2026). However, our results indicate that antagonists, like pathogens and insect herbivores, could limit the population growth of this expanding plant species under future climate conditions.

*P. lanceolata* had a significantly increased population growth in the combined herbivore and pathogen removal treatment (IP) under ambient climate conditions and benefited from insect (I) and IP reduction under future climate conditions. Compared to other species that only suffer from generalist herbivores, *P. lanceolata* hosts both generalist and specialist herbivores (Bowers, 1984; Harvey et al., 2005) and is known to be strongly affected by fungal pathogens (Dudycha and Roach, 2003; Laine, 2003; Xu et al., 2024). Comparably, the reduction of pathogens (P) does not affect the population growth of *P. lanceolata* positively compared to C and leads to a decrease in population size under future climate conditions. This could be due to other plant species benefiting strongly from the exclusion of pathogens, which can give them a competitive advantage over the studied species, counteracting positive effects of pathogen reduction (Mordecai, 2011).

The other two study species, *D. carthusianorum* and *T. orientalis,* showed only weak effects to the separate and combined effects of the climate and antagonist treatments. *D. carthusianorum* showed marginally significant differences between the ambient and future climate, only when pathogens were excluded. The lack of response to the herbivore and pathogen reduction treatments might be explained by the little overall pathogen or insect damage observed on *D. carthusianorum* (personal observation). Seed predation by insects has been documented in other studies on this species (Collin et al., 2002) as well as reproductive limitations because of anther smut pathogens (Koupilová et al., 2023). Despite this, our insect reduction treatments did not significantly increase this species population growth rate or its fertility. The lack of responses to the climate treatment might be due to *D. carthusianorum* having high drought tolerance (Ellenberg et al., 1991). *T. orientalis* exhibited population declines across all treatment combinations, and no differences in population growth rate across treatments. We expected to see a positive effect of the insect reduction treatments, as we observed leaf damage and seed predation by insects (personal observations). However, the low sample size for this species limited our statistical power and highlights the challenge of studying rare and declining species (MacKenzie et al., 2005).

### Sensitivity of vital rates and contribution of treatments to changes in population growth rates

Interestingly, consistently across all four species we found that their population growth rates were highly sensitive to changes in vital rates associated with reproduction (see also Andrzejak et al. 2024, Lemmer et al. 2019). Climate change is known to reduce seed output due to changes in water availability for other species (Gambín and Borrás, 2010). Our results for *B. erectus* show that reductions in seed production under future climate conditions contributed disproportionately to the observed reduction in population growth rate between ambient and future climate treatments. Long-lived species typically exhibit higher sensitivity in λ to survival rather than reproduction (Kuss et al., 2008; Ramula et al., 2008; Weppler et al., 2006). However, species with high λ values also tend to be sensitive to reproductive changes (Franco & Silvertown, 2004; Ramula et al., 2008). Our results indicate that *B. erectus* and *D. carthusianorum*, which had high λ values, were particularly sensitive to reproductive changes, while *T. orientalis* and *P. lanceolata* were more sensitive to changes in growth and reproduction depending on the treatment.

### Future directions

Biotic interactions may shift in response to climate change, potentially leading to the disruption of plant-antagonist relationships (Gellesch et al., 2013; Surówka et al., 2020). While our experiment provides valuable insights into how the joint impact of climate change and local biotic interactions affect plant population dynamics within a single site, its small spatial scale and fixed location mean that highly mobile antagonists (e.g., insects) respond to our treatments through their behaviour rather than through their population dynamics. Thus, we could not observe the full set of potential changes between plants and antagonists that could happen on a larger spatial scale (e.g., through shifts in distributions). In order to overcome this limitation, multi-site exclusion climate change experiments, where plant populations and communities are exposed to climate treatments along with different antagonist reduction treatments could help to further understand future plant-antagonist dynamics.

## Conclusions

Despite the urgent need to understand and mitigate climate-related changes in plant-antagonist interactions (e.g. for conservation of plant species), experimental studies on this topic remain scarce, especially those that consider multiple antagonists like insects and pathogens (Gilman et al., 2010; Pettorelli, 2012; Pressey et al., 2007). Our findings highlight potential interactive and synergistic effects between different antagonist groups on plant populations and show that they can mediate plant responses to climate change. By incorporating demographic responses, we are able to pinpoint the underlying causes of population level changes.

## Author contributions

The study was designed by Lotte Korell (leading), Martin Andrzejak and Tiffany M. Knight (both supporting). Fieldwork was designed by Lotte Korell and Martin Andrzejak (both leading) and carried out by Martin Andrzejak (leading) and Lotte Korell (supporting). Analysis was carried out by Martin Andrzejak (leading) with the support of the other coauthors. The manuscript was written by Martin Andrzejak (leading) and Tiffany M. Knight and Lotte Korell (supporting). All authors commented and improved the manuscript.

## Acknowledgements

This work was supported by the Alexander von Humboldt Foundation (Alexander von Humboldt Professorship held by Tiffany M. Knight) and the Helmholtz Recruitment Initiative of the Helmholtz Association (Tiffany M. Knight). We gratefully acknowledge iDiv, funded by the German Research Foundation (DFG–FZT 118), for its support. The GCEF project was funded by the Helmholtz Association, the German Federal Ministry of Education and Research, the State Ministry of Science and Economy of Saxony-Anhalt, and the State Ministry for Higher Education, Research and the Arts of Saxony.

We sincerely thank the staff of the Bad Lauchstädt Experimental Research Station, in particular Ines Merbach and Konrad Kirsch, for their continued maintenance of the GCEF infrastructure and experimental plots as well as Sigrid Berger for establishing and maintaining the treatments within the GCEF. We are also grateful to Harald Auge, François Buscot, Stefan Klotz, Thomas Reitz, and Martin Schädler for their contributions to the establishment of the GCEF. Field assistance provided by Julia Lemmer, Christian Savona, Simon Bitzan, Georg Küstner, Lina Lüttgert, and Amibeth Thompson is greatly appreciated. We further thank Aldo Compagnoni for analytical support and Harald Auge for insightful comments and discussions that improved this study as well as Amibeth Thompson for providing valuable feedback.

## Conflict of interest statement

The authors declare no conflicts of interest

## Data availability statement

The data and code will be made available upon submission on figshare.

## Appendix

### Figures

**Figure A1.**
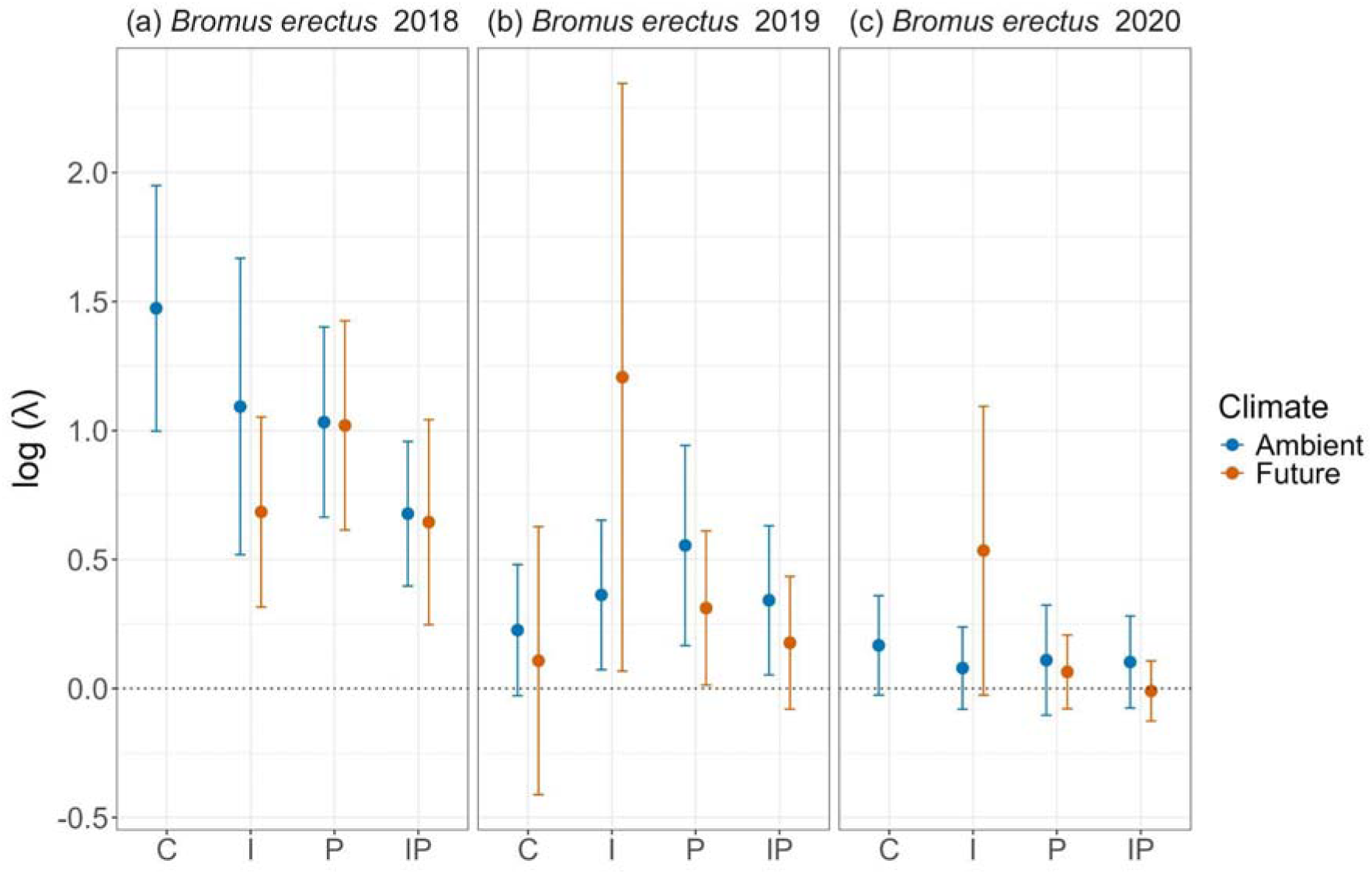
Yearly IPM’s for *Bromus erectus*. Abbreviations for the x axis are as follows: C = control, I = insect, P = pathogen, IP = insect and pathogen. Due to insufficient data some treatment combinations are missing in different years.

**Figure A2.**
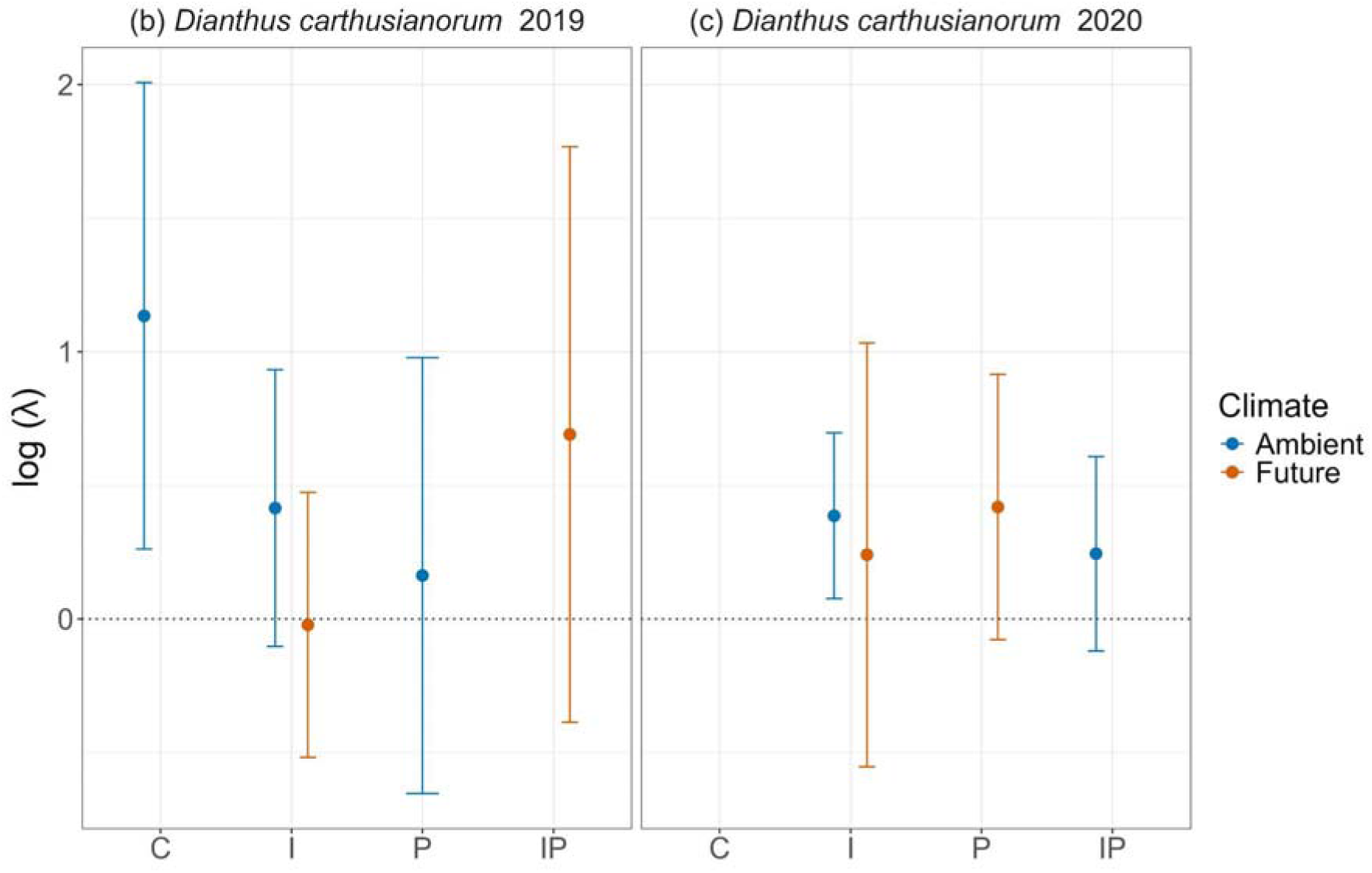
Yearly IPM’s for *Dianthus carthusianorum*. Abbreviations for the x axis are as follows: C = control, I = insect, P = pathogen, IP = insect and pathogen. Due to insufficient data for some vital rates some transitions or data points are missing from the plot.

**Figure A3.**
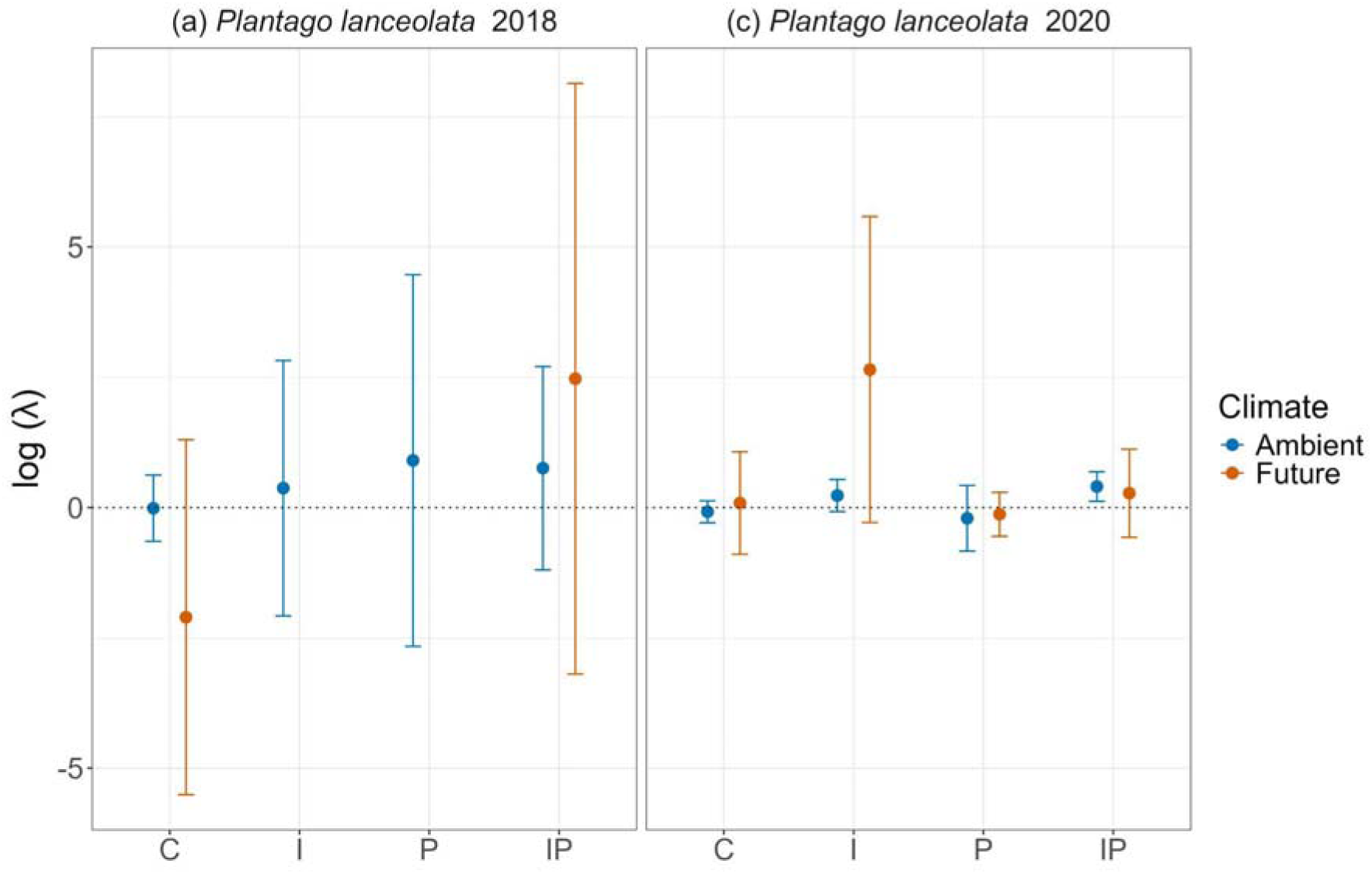
Yearly IPM’s for *Plantago lanceolata*. Abbreviations for the x axis are as follows: C = control, I = insect, P = pathogen, IP = insect and pathogen. Due to insufficient data some transitions or data points are missing from the plot.

**Figure A4.**
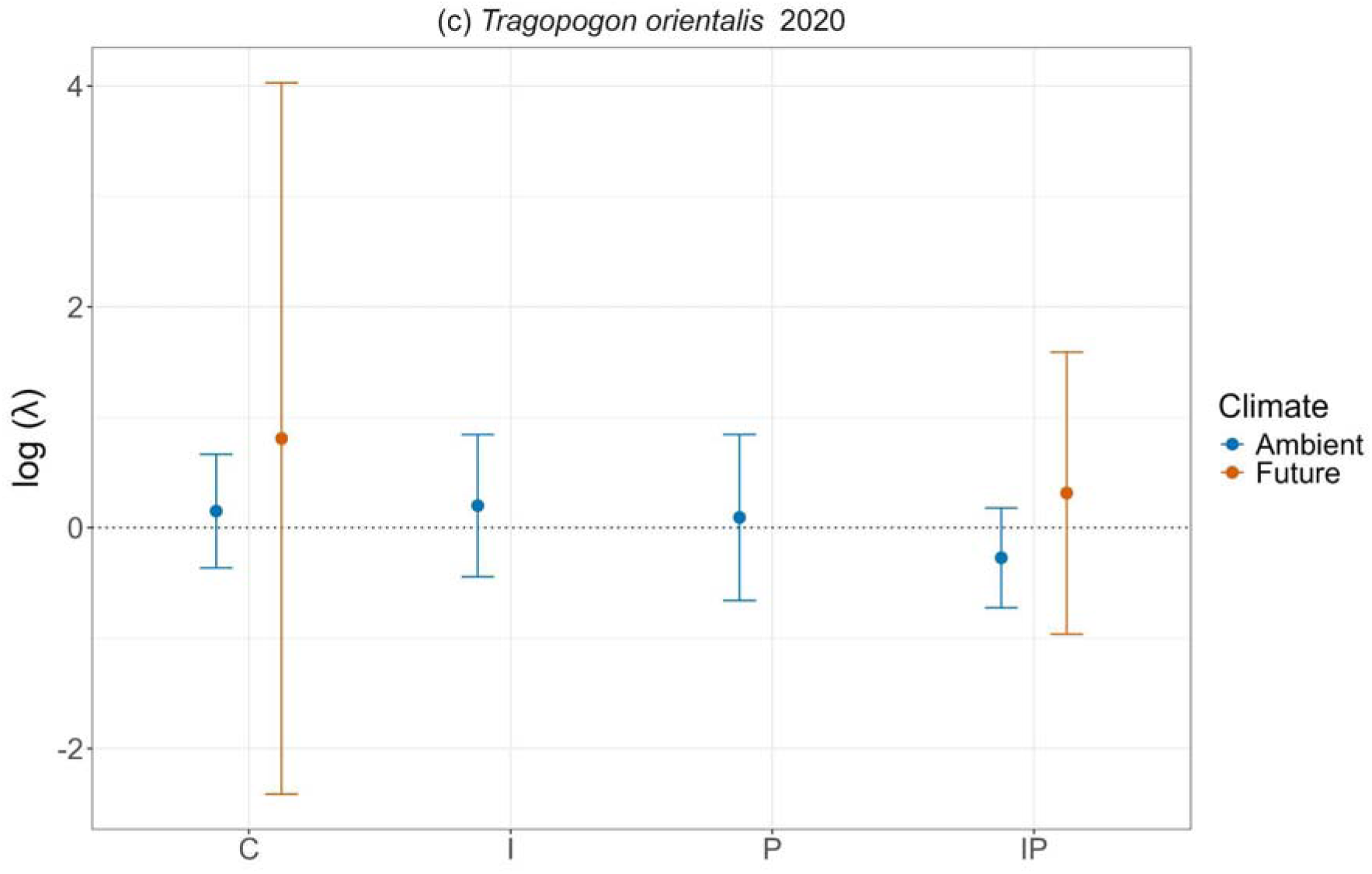
Yearly IPM’s for *Bromus Tragopogon*. Abbreviations for the x axis are as follows: C = control, I = insect, P = pathogen, IP = insect and pathogen. Due to insufficient data some transitions or data points are missing from the plot.

**Figure A5.**
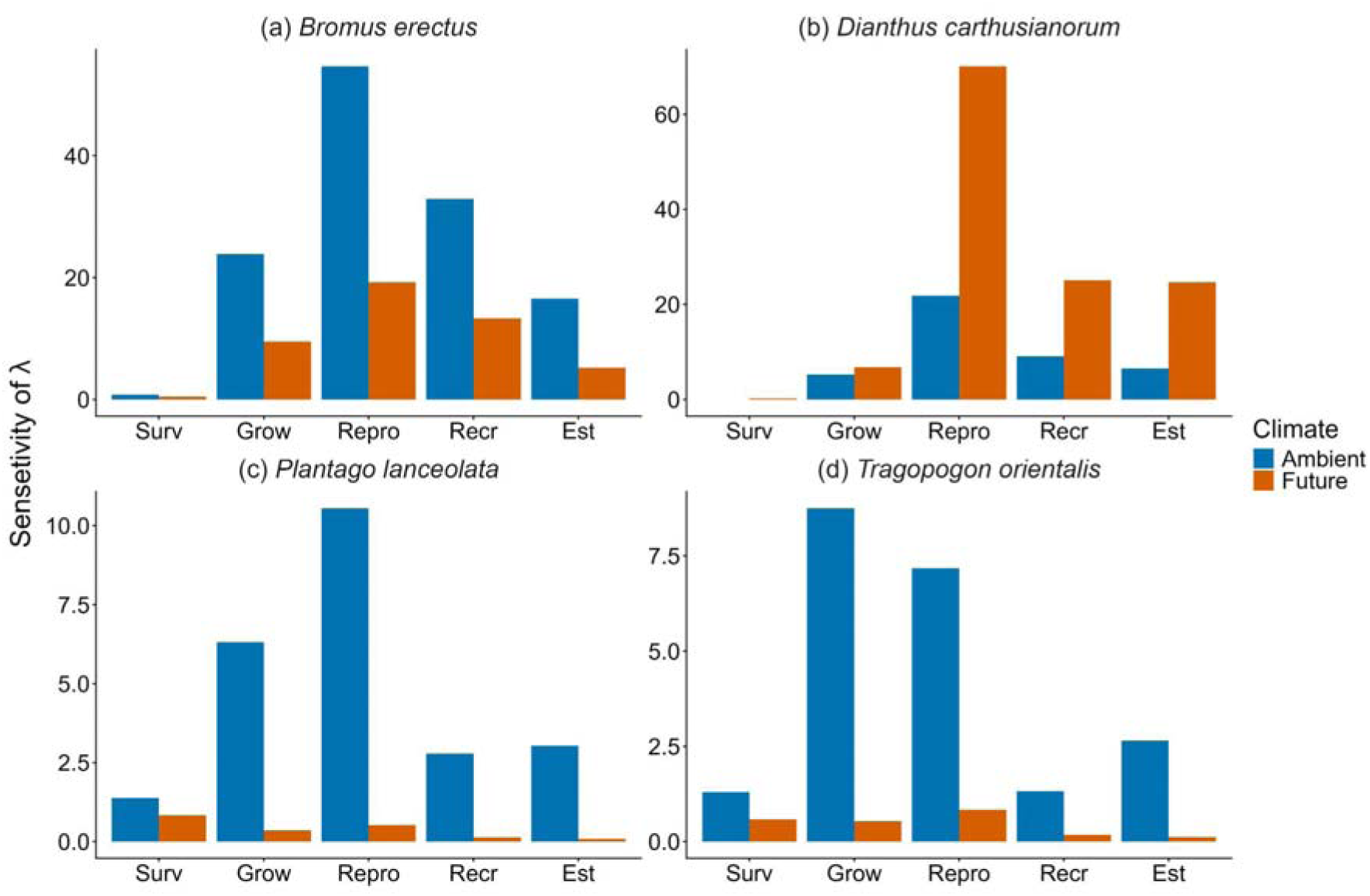
Sensitivity for the two climate treatments. Shown is every vital rate included in the IPM. Note that the scales of the different plots are not on the same scale.

**Figure A6.**
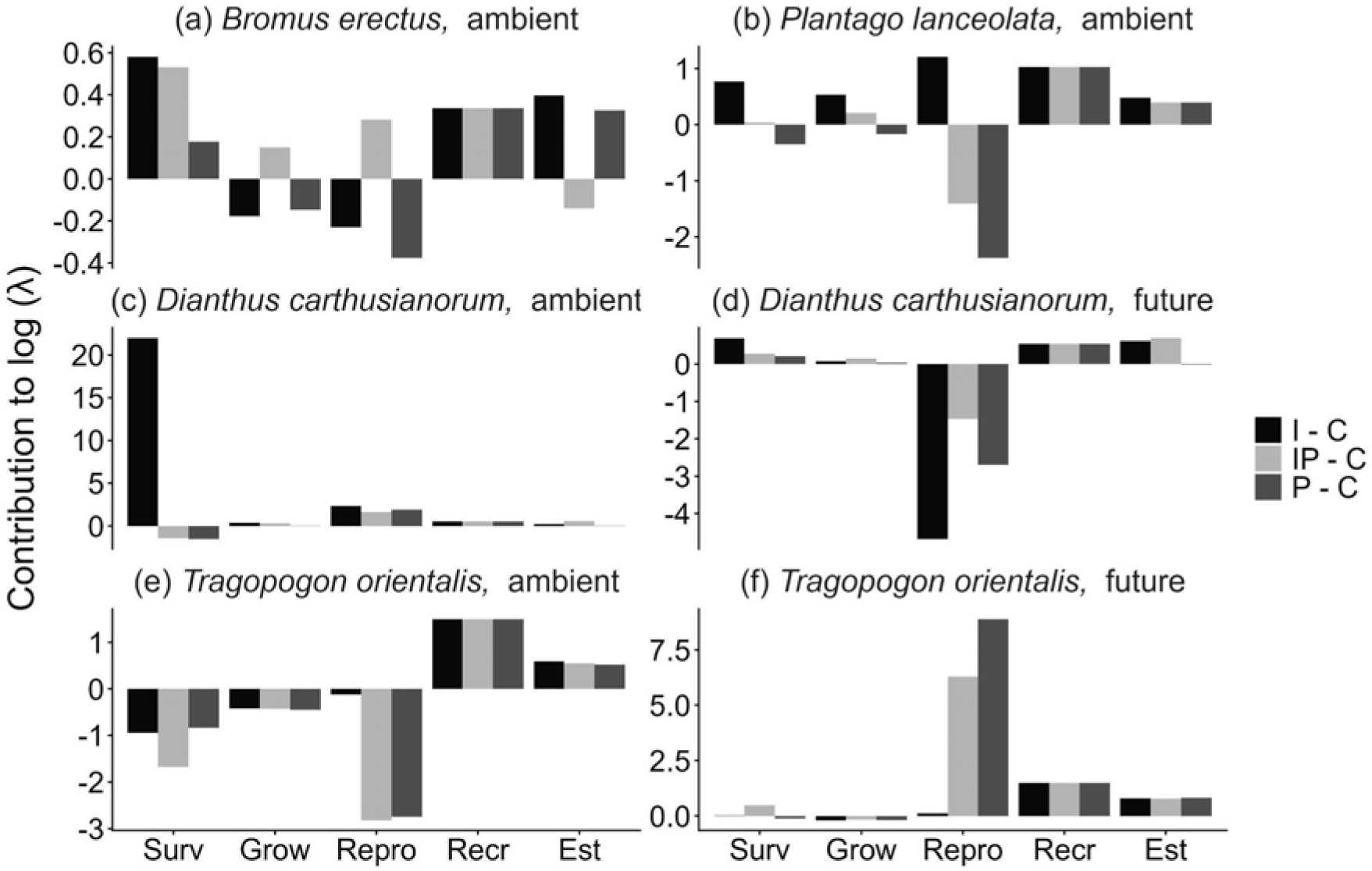
Results of the Life Table Response Experiment (LTRE) for *B. erecuts* and *P. lanceolata* comparing the reduction treatments to the control under ambient climate conditions (a and b). Panels c - f show *D. carthusianorum* and *T. orientalis*. For those two species shown are the results when the reduction treatments are compared to the control under both climate treatments. In the figure legend the treatments are abbreviated: C = control, I = insect, P = pathogen, IP = insect and pathogen. The x axis abbreviations are like this: Est = Establishment, Grow = Growth, Recr = Recruitment, Repro = Reproduction, Surv = Survival. Note that the y-axis are not on the same scale.

**Figure A7.**
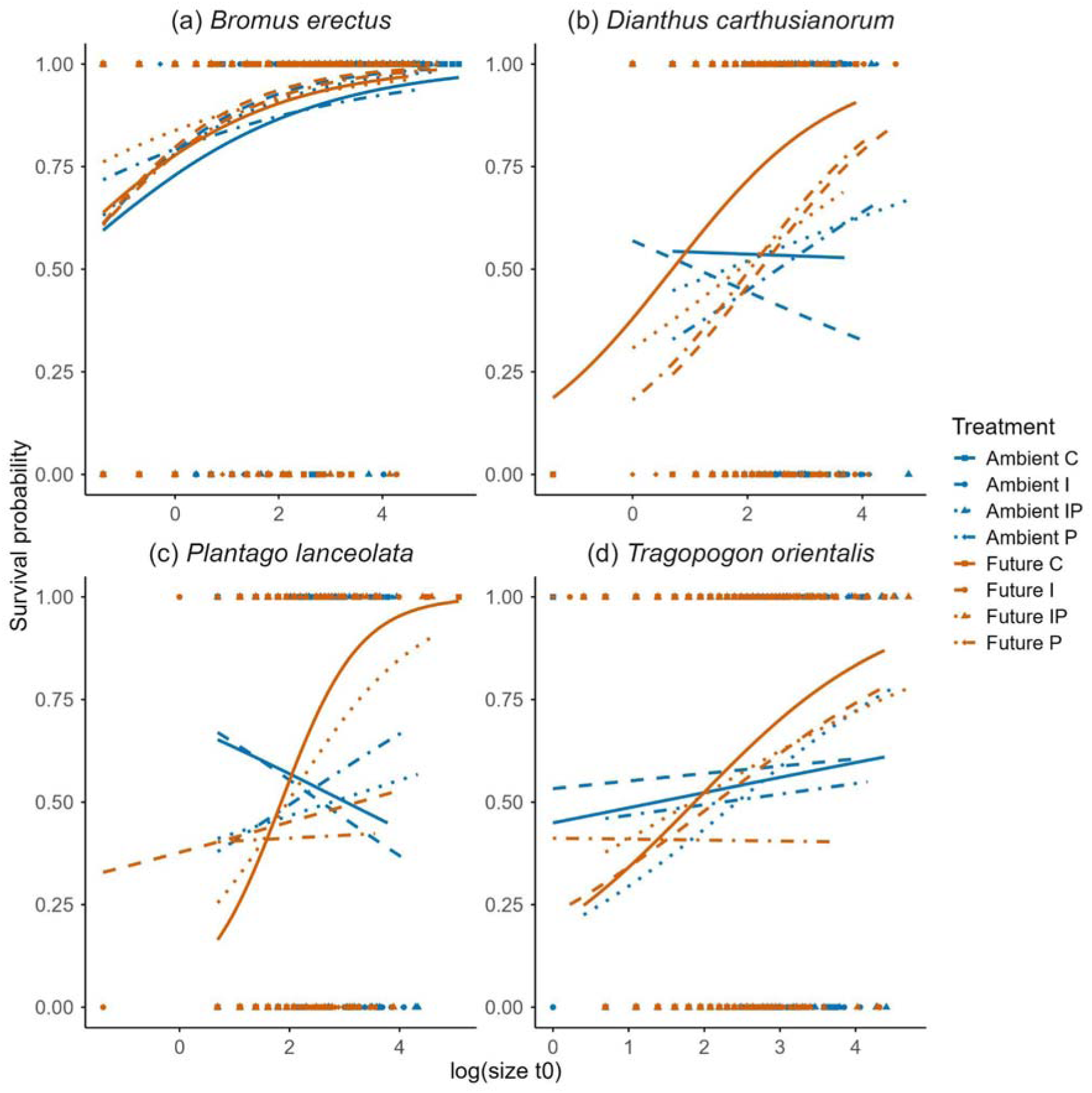
The relation between plant size at t0 and survival for (a) *Bromus erectus* (b) *Dianthus carthusianorum* (c) *Plantago lanceolata* (d) *Tragopogon orientalis*. Different treatment combinations are indicated with different shapes and colors.

**Figure A8.**
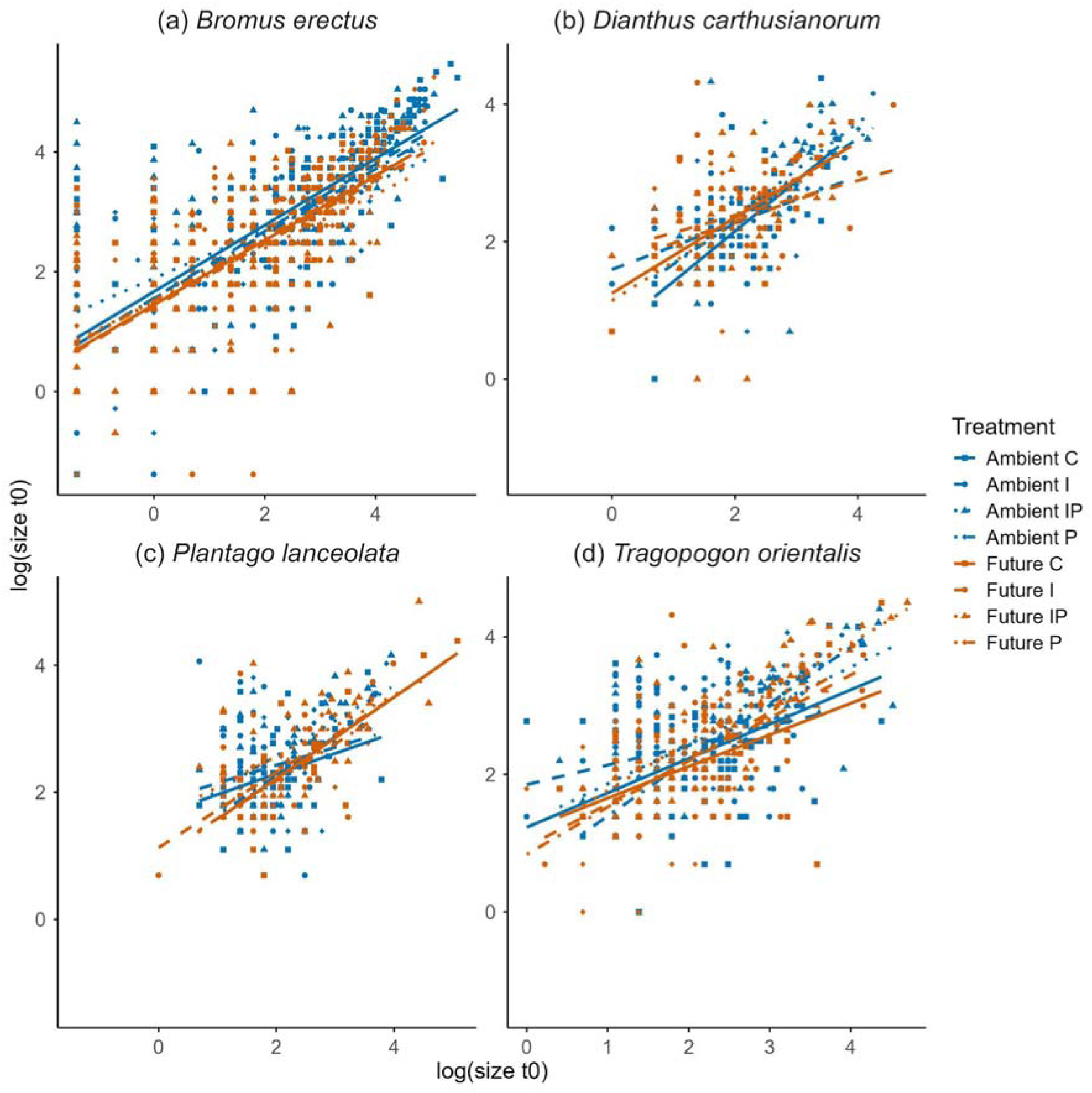
Growth of the different species between size at t0 and size at t1 for (a) *Bromus erectus* (b) *Dinathus carthusianorum* (c) *Plantago lanceolata* (d) *Tragopogon orientalis*. Different treatment combinations are indicated with different colors, shapes and line types.

**Figure A9.**
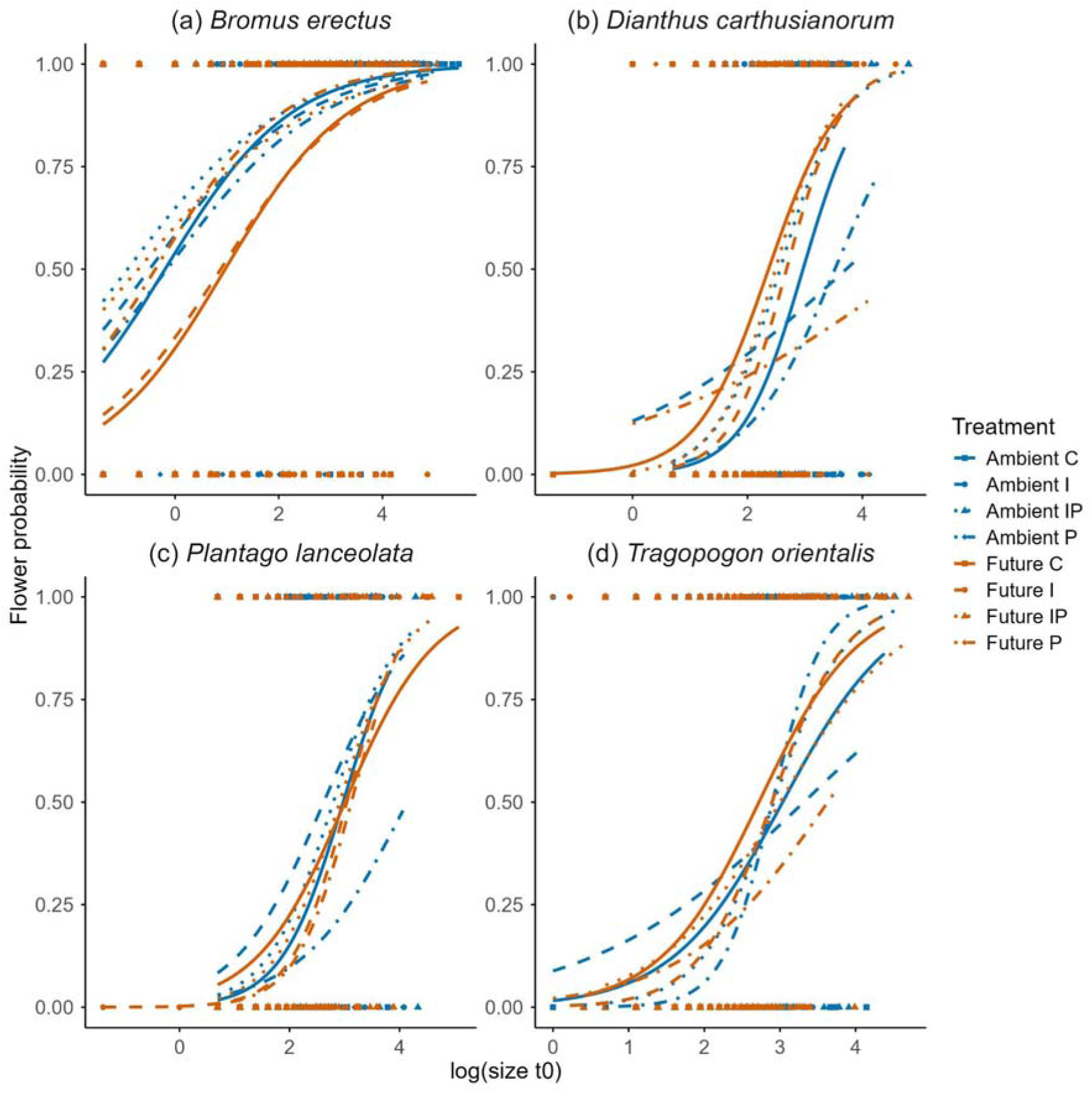
Probability of flowering for each individual of each species at size t0 for (a) *Bromus erectus* (b) *Dianthus carthusianorum* (c) *Plantago lanceolata* (d) *Tragopogon orientalis*. Different treatment combinations are indicated with different colors, shapes and line types.

**Figure A10.**
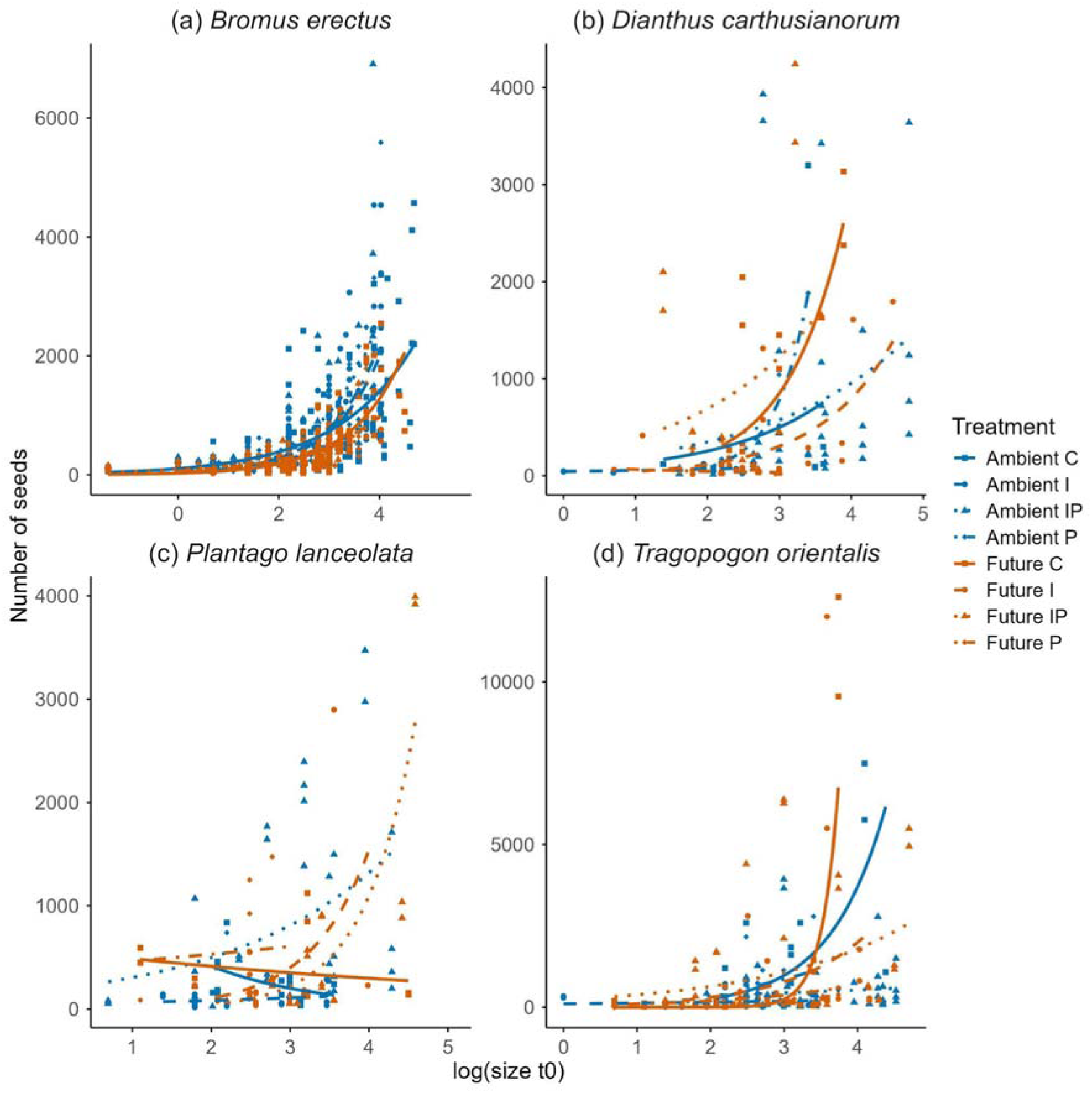
Number of seeds at size t0 for each species (a) *Bromus erectus* (b) *Dianthus carthusianorum* (c) *Plantago lanceolata* (d) *Tragopogon orientalis*. Different treatment combinations are indicated with different colors, shapes and line types.

**Figure A11.**
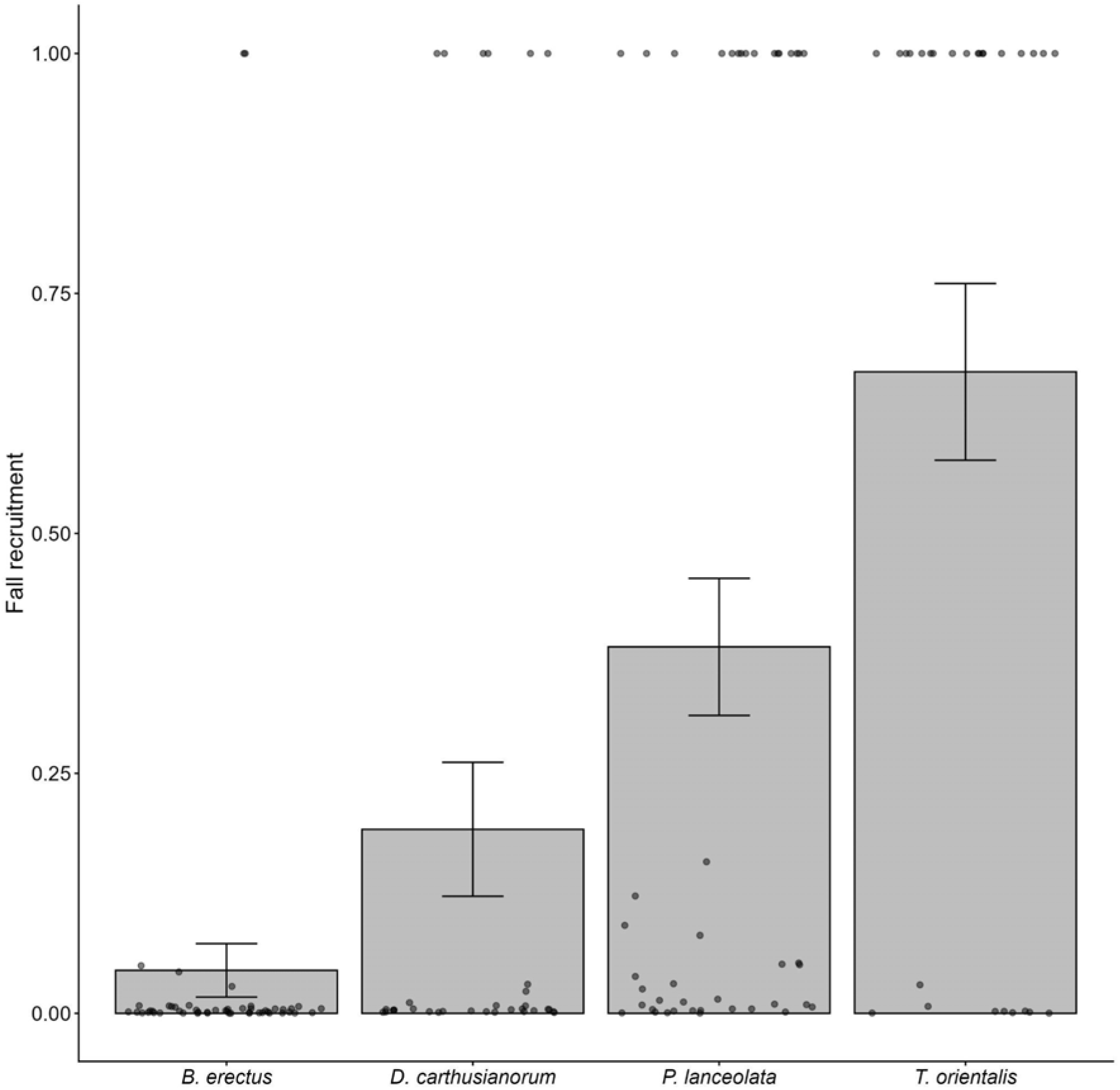
Proportion of seeds that become seedlings in fall. Shown are the four species on the x-axis. Due to little differences between treatments, we pooled the data for this life stage.

**Figure A12.**
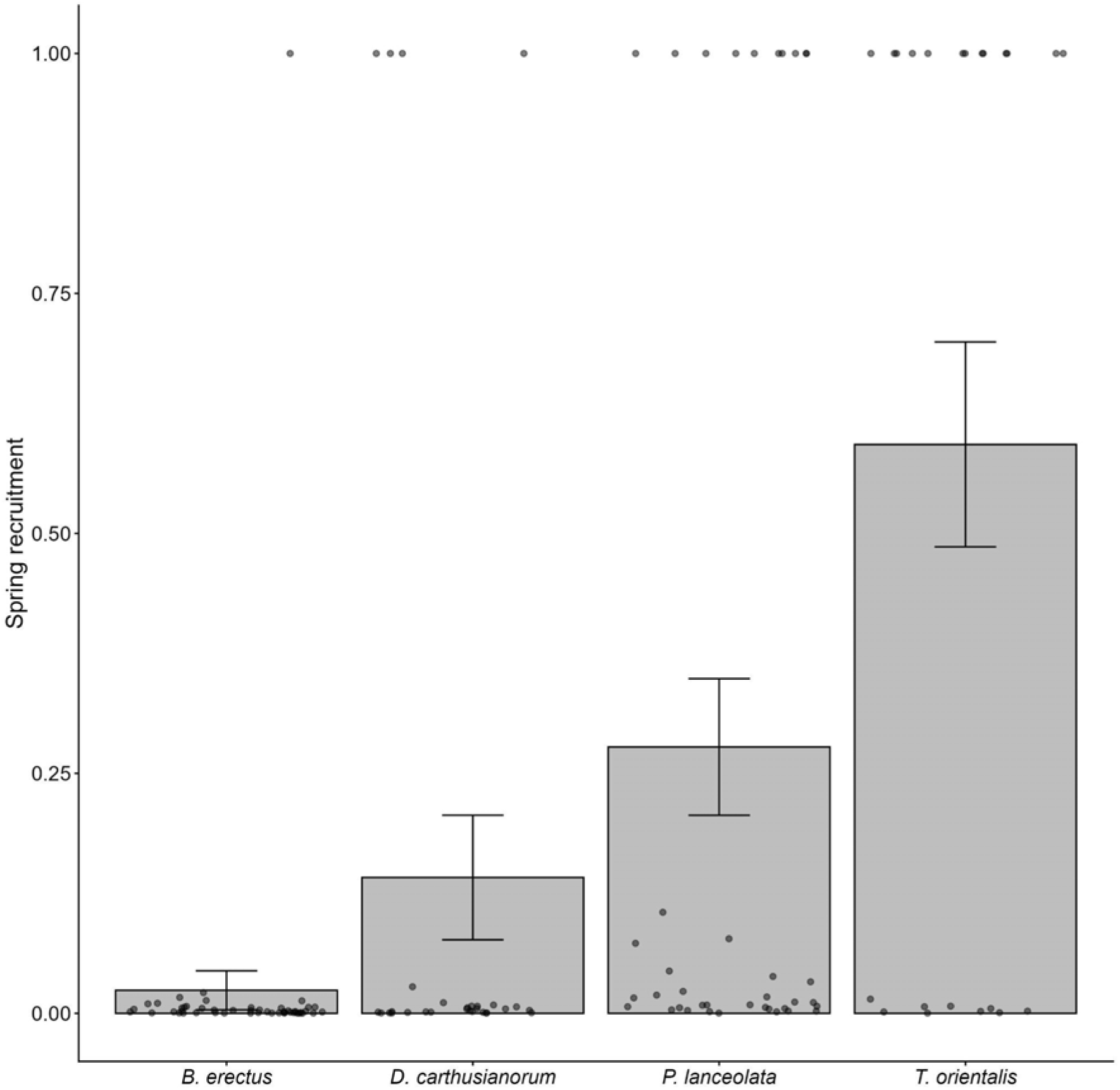
Proportion of seeds that become a seedling in spring. Shown are the four species on the x-axis. Due to little differences between treatments, we pooled the data for this life stage.

**Figure A13.**
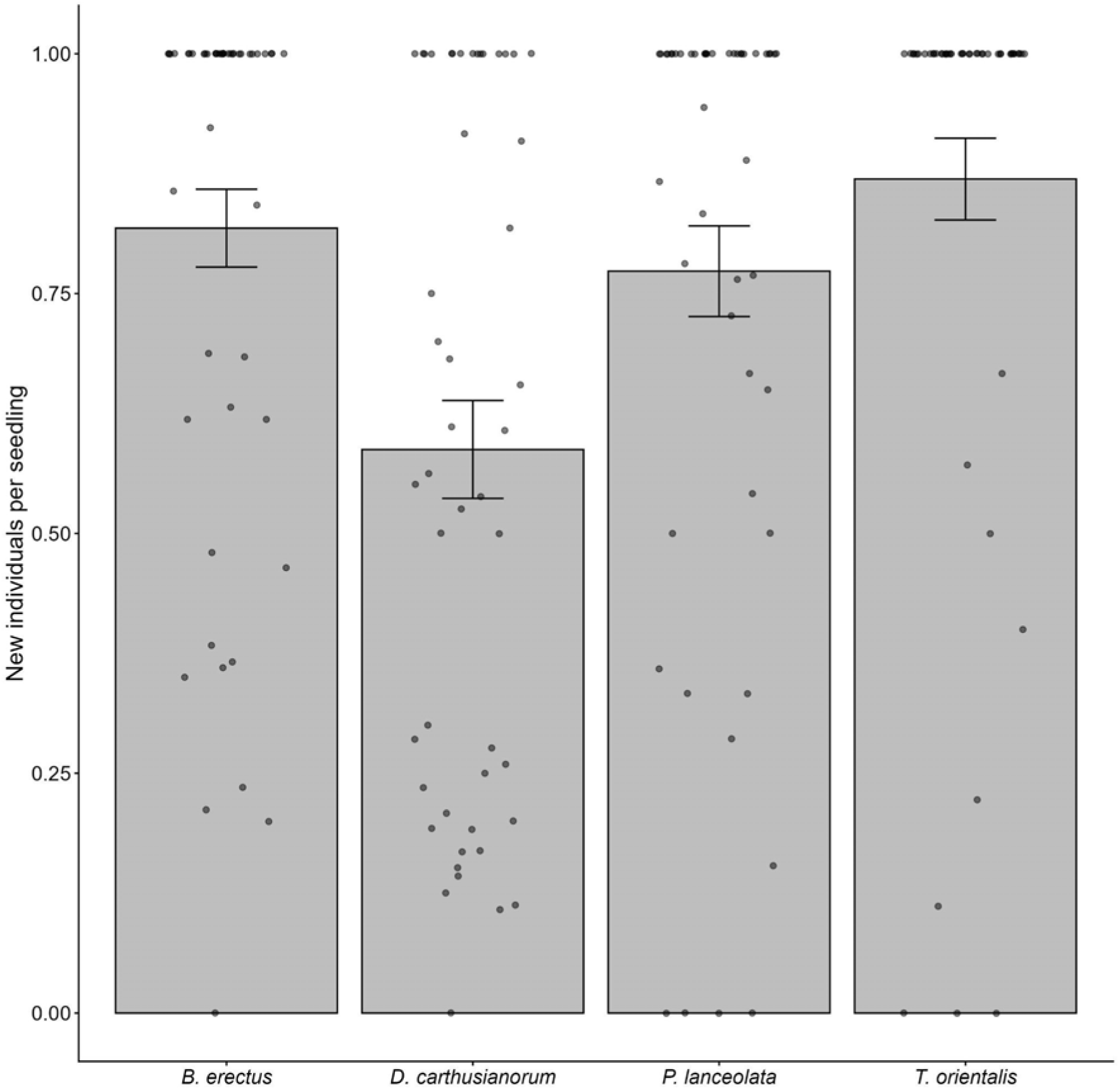
Proportion of seedlings that grow into new adults.

**Figure A14.**
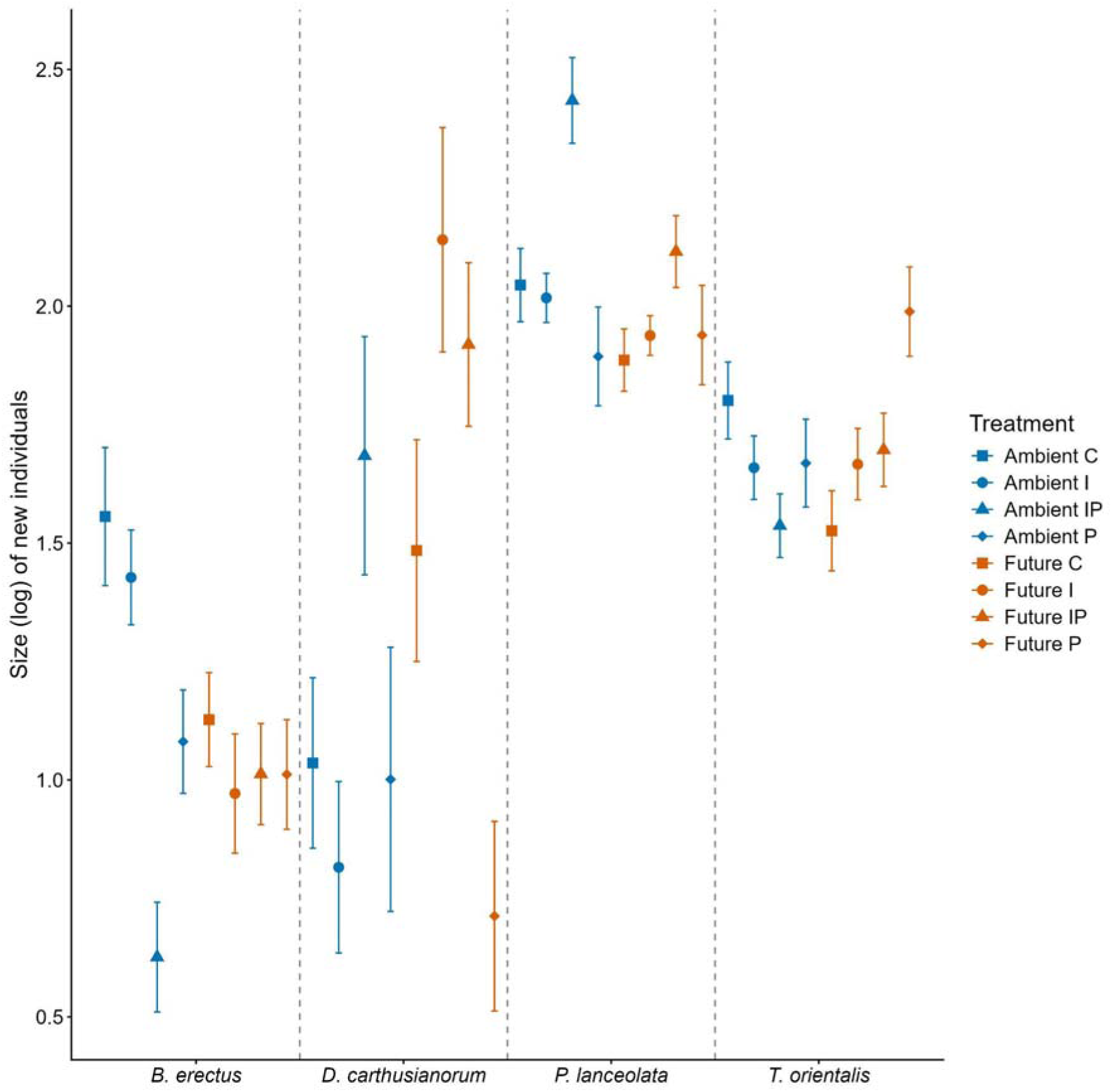
Size distribution of new individuals for each species between each treatment. Blue indicates ambient climate and orange indicates future climates. The different treatment combinations are indicated by different symbols.

**Table A1.**
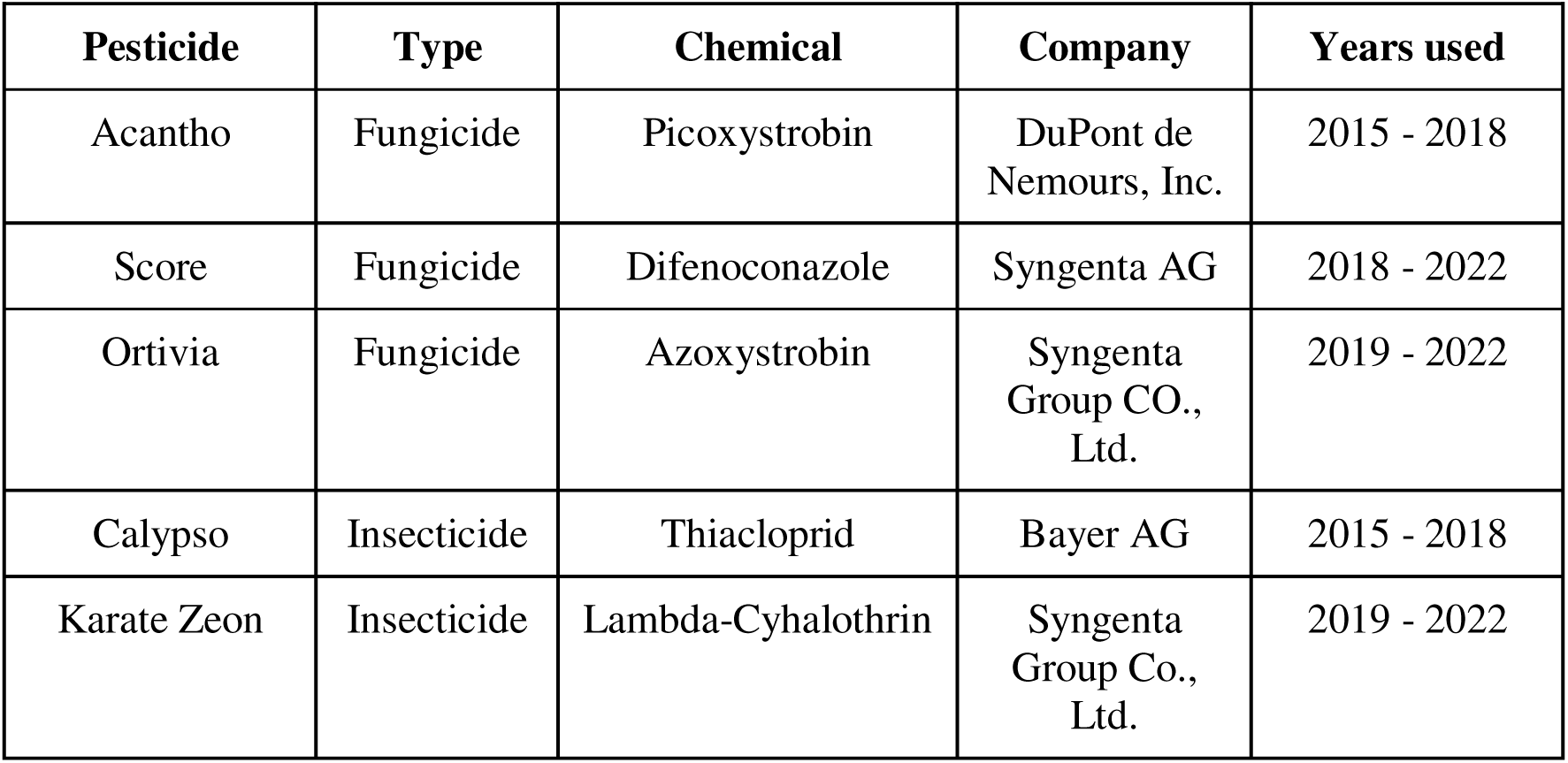
Shown are the chemicals that were applied to the reduction plots and which years they were used. Additional information are the type of pesticide, the active ingredient and the companies which provided the pesticides.

| **Pesticide** | **Type** | **Chemical** | **Company** | **Years used** |
| --- | --- | --- | --- | --- |
| Acantho | Fungicide | Picoxystrobin | DuPont de Nemours, Inc. | 2015 - 2018 |
| Score | Fungicide | Difenoconazole | Syngenta AG | 2018 - 2022 |
| Ortivia | Fungicide | Azoxystrobin | Syngenta Group CO., Ltd. | 2019 - 2022 |
| Calypso | Insecticide | Thiacloprid | Bayer AG | 2015 - 2018 |
| Karate Zeon | Insecticide | Lambda-Cyhalothrin | Syngenta Group Co., Ltd. | 2019 - 2022 |

## Notes

### Competing Interest Statement

The authors have declared no competing interest.

